# Membrane lectins enhance SARS-CoV-2 infection and influence the neutralizing activity of different classes of antibodies

**DOI:** 10.1101/2021.04.03.438258

**Authors:** Florian A. Lempp, Leah Soriaga, Martin Montiel-Ruiz, Fabio Benigni, Julia Noack, Young-Jun Park, Siro Bianchi, Alexandra C. Walls, John E. Bowen, Jiayi Zhou, Hannah Kaiser, Maria Agostini, Marcel Meury, Exequiel Dellota, Stefano Jaconi, Elisabetta Cameroni, Herbert W. Virgin, Antonio Lanzavecchia, David Veesler, Lisa Purcell, Amalio Telenti, Davide Corti

**Affiliations:** Vir Biotechnology, San Francisco, CA 94158, USA; Humabs Biomed SA, a subsidiary of Vir Biotechnology, 6500 Bellinzona, Switzerland; Department of Biochemistry, University of Washington, Seattle, WA 98195, USA; Department of Pathology and Immunology, Washington University School of Medicine, Saint Louis MO 63110, USA; Department of Internal Medicine, UT Southwestern Medical Center, Dallas, TX 75390, USA; Vir Biotechnology, St Louis, MO, 63110, USA

**Keywords:** SARS-CoV-2, COVID-19, antibody, vaccine, neutralising antibodies, mutation, lectin receptors, auxiliary-receptors, sialic acid

## Abstract

Investigating the mechanisms of SARS-CoV-2 cellular infection is key to better understand COVID-19 immunity and pathogenesis. Infection, which involves both cell attachment and membrane fusion, relies on the ACE2 receptor that is paradoxically found at low levels in the respiratory tract, suggesting that additional mechanisms facilitating infection may exist. Here we show that C-type lectin receptors, DC-SIGN, L-SIGN and the sialic acid-binding Ig-like lectin 1 (SIGLEC1) function as auxiliary receptors by enhancing ACE2-mediated infection and modulating the neutralizing activity of different classes of spike-specific antibodies. Antibodies to the N-terminal domain (NTD) or to the conserved proteoglycan site at the base of the Receptor Binding Domain (RBD), while poorly neutralizing infection of ACE2 over-expressing cells, effectively block lectin-facilitated infection. Conversely, antibodies to the Receptor Binding Motif (RBM), while potently neutralizing infection of ACE2 over-expressing cells, poorly neutralize infection of cells expressing DC-SIGN or L-SIGN and trigger fusogenic rearrangement of the spike promoting cell-to-cell fusion. Collectively, these findings identify a lectin-dependent pathway that enhances ACE2-dependent infection by SARS-CoV-2 and reveal distinct mechanisms of neutralization by different classes of spike-specific antibodies.

## Introduction

SARS-CoV-2 infects target cells via the spike glycoprotein (S) that is organized as a homotrimer wherein each monomer is comprised of S1 and S2 subunits^1,2^. The infectious process includes binding to cells, triggering of S conformational changes and then fusion of the viral envelope with the target cell membrane. The S1 subunit comprises the N-terminal domain (NTD) and the receptor binding domain (RBD), the latter interacting with ACE2 through a region defined as the receptor binding motif (RBM). Antibodies against the RBD contribute to the majority of the neutralizing activity in polyclonal serum antibodies^3,4^, potently neutralize SARS-CoV-2 in vitro^5,6^ and have shown efficacy in clinical trials for prophylaxis and early therapy of COVID-19^7,8^.

The search for SARS-CoV-2 neutralizing antibodies has been facilitated by the use of target cells over-expressing the ACE2 receptor^9^. However, while ACE2 is highly expressed in several tissues including the intestine and kidney, its expression in the respiratory tract is limited, with low levels found in only a limited number of type-II alveolar basal, goblet and mucous cells^10–12^. The paradox of low ACE2 levels in the lung and infection in other tissues leading to extrapulmonary complications^13^, raises the possibility that additional receptors may contribute to viral infection and dissemination, such as DC-SIGN (CD209), L-SIGN (CD209L/CLEC4M), neuropilin-1 (NRP-1), basigin (CD147) and heparan sulfate^14–18^. It remains to be established whether these molecules may act as alternative receptors for viral entry, as co-receptors or as auxiliary receptors that tether viral particles enhancing their interaction with ACE2.

In this study, we identify DC-SIGN, L-SIGN and SIGLEC1 as auxiliary receptors that enhance ACE2-dependent infection and demonstrate different mechanisms of neutralization by antibodies targeting RBM and non-RBM sites in the presence or absence of lectins.

## Results

### DC-SIGN, L-SIGN and SIGLEC1 act as auxiliary receptors for ACE2-dependent SARS-CoV-2 infection

To develop an assay for identification of accessory receptors of SARS-CoV-2 infection, we took advantage of HEK293T cells that express low endogenous levels of ACE2. HEK293T were transfected with vectors encoding ACE2 or 13 selected lectins and published receptor candidates prior to infection with VSV-SARS-CoV-2. As expected, untransfected HEK293T cells were only weakly permissive to infection, and ACE2 overexpression led to a dramatic increased pseudovirus entry. Interestingly, increased infectivity was also observed in HEK293T cells following transfection with C-type lectins DC-SIGN and L-SIGN that were previously reported as entry receptors^14,15,19^, as well as with SIGLEC1, which was not previously shown to mediate SARS-CoV-2 entry (**Fig. 1a**). In contrast, NRP-1 and CD147 did not enhance_SARS-CoV-2 infection in these conditions although they were previously suggested to act as entry receptors^16,17^. The infection-enhancing activity of the three lectins was also observed with authentic SARS-CoV-2 and stably transduced HEK293T cells (**Fig. 1b-c** and **Extended Data Fig. 1**). A SIGLEC1 blocking antibody inhibited in a dose-dependent fashion the infection of SIGLEC1 expressing HEK293T, supporting the role of this molecule as a new SARS-CoV-2 entry factor (**Fig. 1d**).

**Fig. 1.**
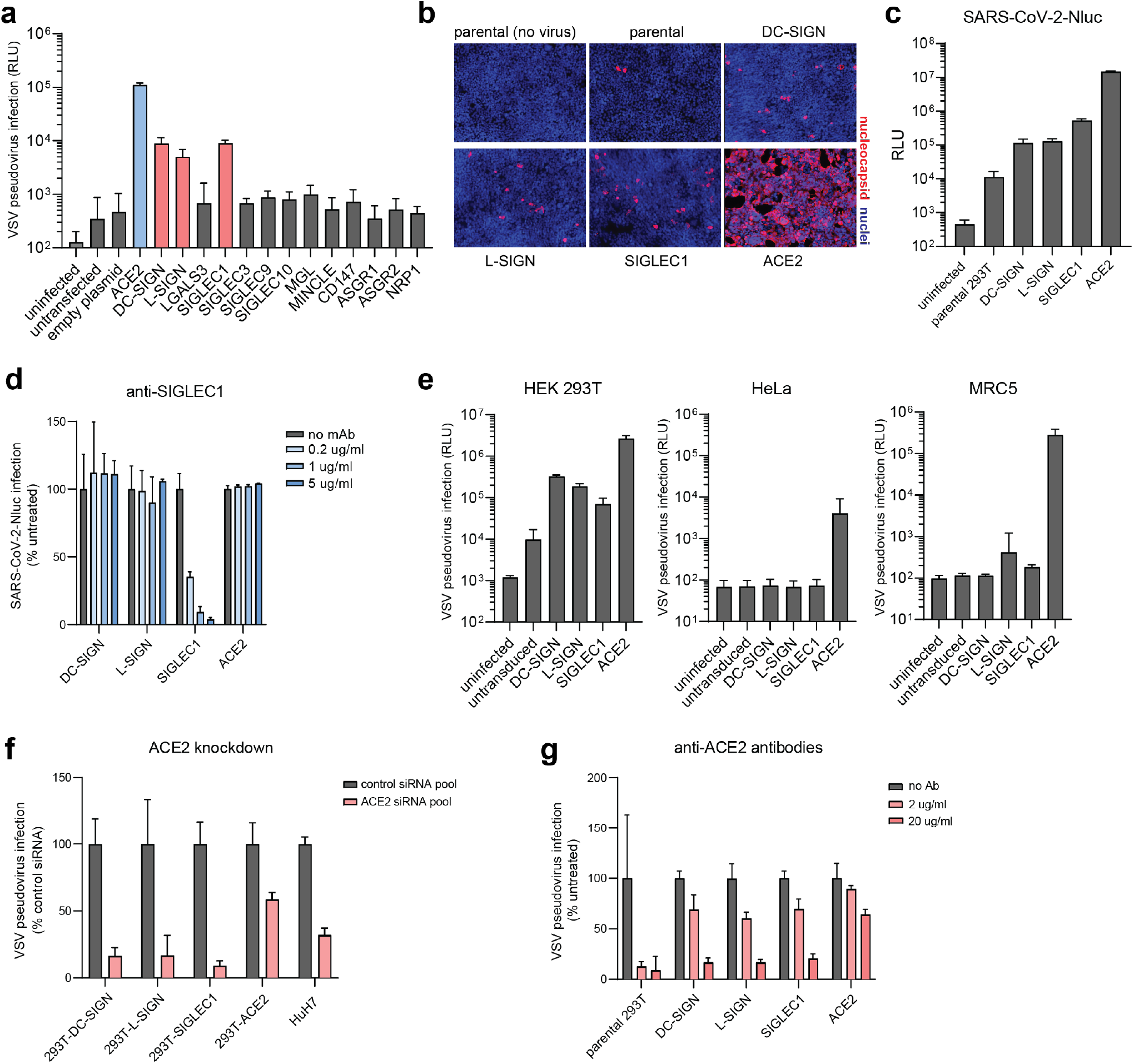
DC-SIGN, L-SIGN and SIGLEC1 function as auxiliary receptors for SARS-CoV-2 infection. **a,** VSV-SARS-CoV-2 pseudovirus infection of HEK293T cells transfected to over-express ACE2 or a panel of selected lectins and published receptor candidates. **b,** Stable HEK293T cell lines overexpressing DC-SIGN, L-SIGN, SIGLEC1 or ACE2 were infected with authentic SARS-CoV-2 (MOI 0.1), fixed and immunostained at 24 hours for the SARS-CoV-2 nucleocapsid protein (red). **c,** HEK293T stable cell lines were infected with SARS-CoV-2-Nluc and luciferase levels were quantified at 24 hours. **d,** Stable cell lines were incubated with different concentrations of anti-SIGLEC1 mAb (clone 7-239) and infected with SARS-CoV-2-Nluc. **e,** HEK293T, HeLa and MRC5 cells were transiently transduced to overexpress DC-SIGN, L-SIGN, SIGLEC1 or ACE2 and infected with VSV-SARS-CoV-2 pseudovirus. **f,** Stable cell lines were treated with ACE2 siRNA followed by infection with VSV-SARS-CoV-2 pseudovirus four days post transfection. **g,** Stable cell lines were incubated with different concentrations of anti-ACE2 goat polyclonal antibodies and infected with VSV-SARS-CoV-2 pseudovirus.

The ectopic expression of DC-SIGN, L-SIGN and SIGLEC1 did not support infection of ACE2 negative cells, such as HeLa or MRC-5 cells (**Fig. 1e**), indicating that these lectins do not act as entry receptors. The requirement of ACE2 for viral infection of lectin-expressing cells was also demonstrated using ACE2 blocking antibodies or ACE2-siRNA (**Fig. 1f-g**). Of note, the B.1.1.7 UK variant of concern (VOC) which is currently the most prevalent circulating virus in Europe retained the capacity to use DC-SIGN, L-SIGN and SIGLEC1 as auxiliary receptors (**Extended Data Fig. 1c**).

Collectively, these data reveal a lectin-facilitated pathway of infection that is evident on cells expressing low levels of ACE2, supporting the notion that SARS-CoV-2 may use these lectins as auxiliary receptors for ACE2-mediated entry.

### Auxiliary receptors can facilitate trans-infection of ACE2^+^ cells

The above data suggest that lectin receptors may act as auxiliary receptors that tether viral particles to the cell membrane facilitating interaction with ACE2. This could take place in *cis* (on the same cell) or in *trans* effectively promoting cell-to-cell trans-infection, as reported for HIV-1^20^. To address whether ACE2 and lectins can be found on the same cells, we interrogated the lung cell atlas^21^ and identified the cell types that express these receptors (**Fig. 2a**). DC-SIGN (CD209) is expressed most prominently on IGSF21^+^ dendritic cells, L-SIGN (CLEC4M; CD209L) has a limited expression on vascular structures and SIGLEC1 (CD169) is broadly expressed at the surface of alveolar macrophages, dendritic cells and monocytes. As previously noted, ACE2 expression is limited to subsets of alveolar epithelial type-2, basal and goblet cells. Next, we mined the recently released single-cell transcriptomics data on 3,085 lung epithelial and immune cells obtained from bronchoalveolar lavage fluid or sputum of 8 individuals that suffered from severe COVID-19^22^. The distribution of viral RNA per cell, expressed as log counts per million (logCPM), varied across annotated cell types. Specifically, the content of viral RNA in macrophages is greater relative to secretory cells (2-sided K-S statistic and p-value, D = 0.32092, p-value < 2.2e-16) (**Fig. 2b**). SIGLEC1 was expressed in 41.4% (459/1107 cells) of SARS-CoV-2^+^ macrophages, whereas ACE2 expression was negligible in these cells **(Fig. 2c**). Conversely, ACE2 expression was found in 10.6% (60/565 cells) of SARS-CoV-2^+^ secretory cells, whereas SIGLEC1 expression was negligible **(Fig. 2c**). Plotting SIGLEC1, DC-SIGN and L-SIGN expression as a function of SARS-CoV-2 viral load revealed a strong positive correlation for SIGLEC1 in macrophages (**Fig. 2c**). We confirmed this association in a separate transcriptomics dataset of 1,072 SARS-CoV-2^+^ bronchoalveolar lavage fluid cells from individuals with severe COVID-19^23^. We inspected the available sequenced reads from this dataset to assess the nature of viral RNA in SARS-CoV-2^+^ bronchoalveolar lavage cells. Reads which supported a junction between the 5’ leader sequence and the transcription regulatory sequence (TRS) preceding open reading frames for viral genes were counted as evidence of subgenomic mRNA, a surrogate readout for viral replication. Such reads constituted a small fraction of TRS-containing viral reads, ranging from undetectable to 3.4%, suggesting that minimal replication was occurring in this cell population largely comprised of macrophages and other non-epithelial cell types.

**Fig. 2.**
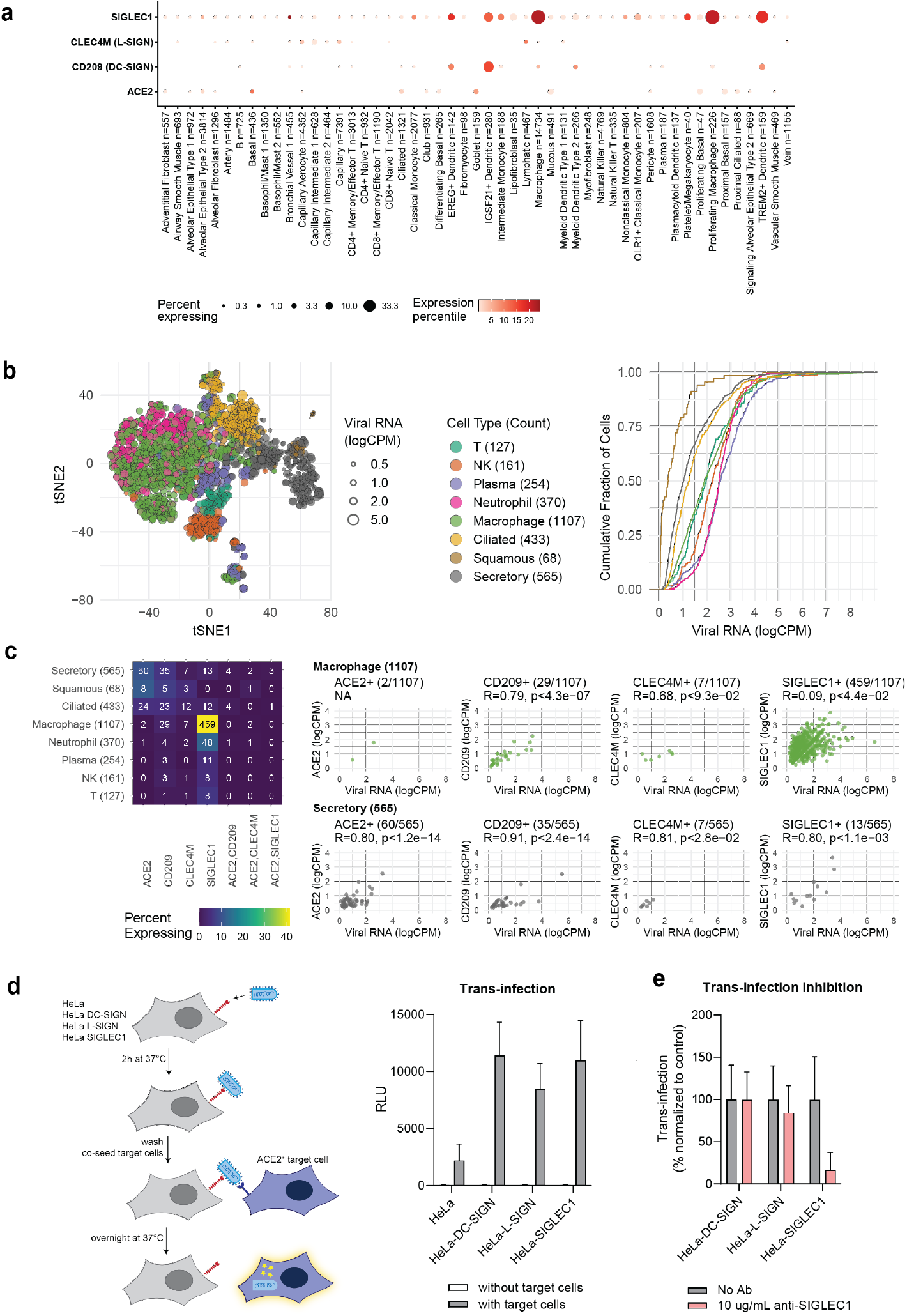
Expression of auxiliary receptors in infected tissues and their role in mediating trans-infection in vitro. **a,** Distribution and expression of ACE2, DC-SIGN, L-SIGN, and SIGLEC1 in the human lung cell atlas. **b,** Major cell types with detectable SARS-CoV-2 genome in bronchoalverolar lavage fluid and sputum of severe COVID-19 patients. Left panel shows a t-SNE embedding of single-cell gene expression profiles coloured by cell type and sized by viral load (logCPM); right panel, distribution plots by annotated cell type denote the cumulative fraction of cells (y-axis) with detected viral RNA per cell up to the corresponding logCPM value (x-axis). **c,** Left panel shows a heatmap matrix of counts for cells with detected transcripts for receptor gene(s) on x-axis by SARS-CoV-2^+^ cell type on y-axis (total n=3,085 cells from 8 subjects in Ren et al.^20^); right panel, correlation of receptor transcript counts with SARS-CoV-2 RNA counts in macrophages and in secretory cells. Correlation is based on counts (before log transformation), from Ren et al.^22^. **d,** Trans-infection: HeLa cells transduced with DC-SIGN, L-SIGN or SIGLEC1 were incubated with VSV-SARS-CoV-2, extensively washed and co-cultured with Vero-E6-TMPRSS2 susceptible target cells. Shown is RLU in the presence or absence of target cells. **e,** Trans-infection performed as in (d). VSV-SARS-CoV-2 viral adsorption was performed in the presence or absence of an anti-SIGLEC1 blocking antibody.

The above results suggest limited cooperation of ACE2 and SIGLEC1 in *cis* because these receptors are rarely expressed in the same cell. However, the data support the possibility of *trans*-infection from SIGLEC1^+^ macrophages to ACE2^+^ cells. To test this hypothesis, we developed an in vitro model where single-round VSV-SARS-CoV-2 was adsorbed on ACE2-negative HeLa cells either untransduced or transduced with DC-SIGN, L-SIGN or SIGLEC1. Under these conditions, lectin-transduced HeLa cells showed enhanced capacity to promote VSV-SARS-CoV-2 trans-infection of susceptible Vero-E6-TMPRSS2 target cells (**Fig. 2d**), and SIGLEC1-mediated trans-infection was inhibited by SIGLEC1-blocking antibodies (**Fig. 2e**). Collectively, these results are consistent with the possible role of lectins in the enhancement of ACE2-mediated trans-infection leading to dissemination of SARS-CoV-2.

### Over-expression of ACE2 reduces neutralization of SARS-CoV-2 by mAbs that do not block ACE2 attachment

Cell line selection and the level of ACE2 expression are important variables in assessing the potency of SARS-CoV-2 neutralizing mAbs. Previous studies suggested that non-RBM mAbs, such as S309 (parent of the clinical stage VIR-7831 antibody) and NTD-specific mAbs showed a reduced and partial neutralizing activity when using target cells over-expressing ACE2^24–26^.

To further investigate how ACE2 and auxiliary receptor expression levels influence neutralizing activity, we compared three mAbs targeting distinct sites on the spike protein: i) S2E12, targeting the RBM site Ia/class 1 on RBD^5^; ii) S309 targeting the conserved proteoglycan site IV/class 3 distal from RBM^27^ and iii) S2X333, targeting site i on NTD^28^ (**Fig. 3a**). These mAbs completely neutralize infection of Vero E6 cells with authentic SARS-CoV-2, albeit with different potencies, and their activity was not influenced by the expression of the TMPRSS2 protease (**Fig. 3b-c and Extended Data Fig. 2a**). To understand the influence of receptor expression on neutralization, we used cell lines expressing ACE2 and TMPRSS2 (endogenously or upon transduction) at levels varying more than 1000-fold as evaluated by qPCR and by flow cytometry staining with RBD or S proteins (**Fig. 3d-e** and **Extended Data Fig. 2b**). Whereas the RBM mAb, S2E12 showed comparable neutralizing activity on all target cells, both S309 and S2X333 showed an impaired neutralizing activity when tested on cells over-expressing ACE2, both in terms of maximal neutralization and potency (**Fig. 3f-g** and **Extended Data Fig. 2c**). This observation was particularly evident for S309 and S2X333 when tested on HEK293T cells over-expressing ACE2. Comparable results were obtained with both VSV-SARS-CoV-2 and authentic SARS-CoV-2-Nluc. Overall, a negative correlation was found between ACE2 levels and neutralization potency for non-RBM mAbs (**Extended Data Fig. 2c).**

**Fig. 3.**
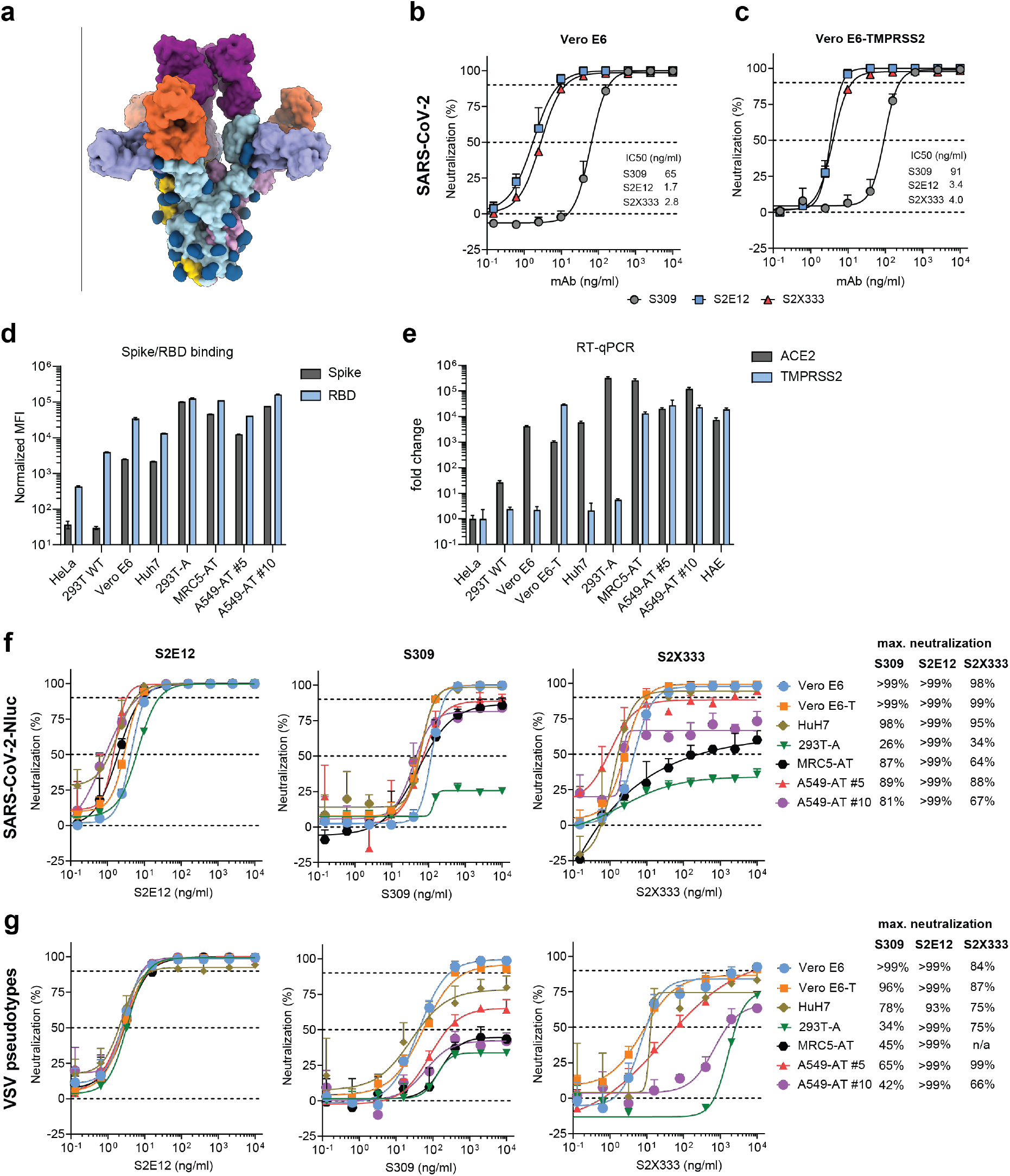
ACE2 over-expression influences neutralizing activity by different classes of anti-spike mAbs. **a**, Surface rendering of a composite model of SARS-CoV-2 S bound to S309 (purple), S2E12 (magenta) and S2X333 (orange)^5, 27, 28^. The three SARS-CoV-2 S protomers are colored light blue, gold and pink whereas N-linked glycans are rendered dark blue. **b-c**, SARS-CoV-2 neutralization with S309, S2E12 and S2X33 on (b) Vero E6 or (c) Vero E6-TMPRSS2 cells. Cells were infected with SARS-CoV-2 (isolate USA-WA1/2020) at MOI 0.01 in the presence of the respective mAbs. Cells were fixed 24h post infection, viral nucleocapsid protein was immunostained and quantified. **d**, Purified, fluorescently-labeled SARS-CoV-2 spike or RBD protein binding to the indicated cell lines was quantified by flow cytometry. “A”: ACE2, “T”: TMPRSS2 **e**, Cellular ACE2 and TMPRSS2 transcripts were quantified by RT-qPCR. **f-g**, A panel of 7 cell lines were infected with SARS-CoV-2-Nluc **f**, or VSV-SARS-CoV-2 pseudovirus **(g)** in the presence of S309, S2E12 or S2X333. Luciferase signal was quantified 24h post infection.

These results demonstrate that the potency of neutralizing mAbs targeting epitopes outside the RBM negatively correlates with ACE2 expression levels on target cells. Consequently, the widespread use of ACE2-overexpressing cells leads to underestimation of the neutralizing activity of non-RBM antibodies, which is well-detected when using cells expressing moderate to low levels of ACE2, possibly corresponding to the physiological ACE2 expression levels.

Given the uncertainty on the most relevant in vitro correlates of protection, we investigated the capacity of hamsterized S309 and S2E12 mAbs to prevent SARS-CoV-2 infection in Syrian hamsters, a relevant animal model that relies on endogenous expression of ACE2^29^. In a prophylactic setting, S309 was highly effective at doses as low as 0.4 mg/kg in terms of reduction of viral RNA and infectious virus levels and histopathological score in the lungs (**Extended Data Fig. 3a**). Furthermore, we did not observe substantial increased efficacy by co-administering S309 with an equal amount of the potent RBM S2E12 mAb (**Extended Data Fig. 3b**). This result is reminiscent of the lack of increased efficacy observed in an early therapy clinical trial where the combination of bamlanivimab and etesevimab mAbs was compared to bamlanivimab alone^30^.

Since S309 induces potent Fc-mediated effector functions in vitro^27^, we set out to determine their contribution in vivo using the hamster model. The S309 mAb harboring the Fc null N297A mutation on the hamster IgG2 Fc failed to bind to hamster monocytes (**Extended Data Fig. 4**), indicating an expected lack of activation of effector functions in vivo. This ‘Fc-silenced’ S309 mAb, however, was similarly protective as wildtype S309 against SARS-CoV-2 challenge of hamsters underscoring that the neutralizing activity of S309 was the primary mechanism of action in this condition (**Extended Data Fig. 5**), which is consistent with the previous finding that in a prophylactic setting in hamsters viral neutralization is the primary mode of action^31^.

Taken together, these data indicate that neutralization assays using cells over-expressing ACE2 under-estimate the neutralizing activity of non-RBM mAbs, which are comparably protective to RBM mAbs in a relevant animal model of infection^32,33^. The significance of this finding is also supported by the recent efficacy data of VIR-7831 in a Phase 3 clinical trial demonstrating 85% protection against hospitalization and death due to COVID-19 in an interim analysis.

### MAb-mediated conformational selection of open RBDs promotes membrane fusion

Infection of permissive cells involves both interactions with ACE2 and auxiliary receptors, as well as fusion of the viral membrane to cellular membranes. Here we investigated how different classes of spike-specific antibodies may interfere with viral fusion events that are involved in viral entry, but also in cell-to-cell fusion, leading to the formation of syncytia in vitro^34^ and of multi nucleate giant cells in human lung from infected individuals^35^. We previously showed that RBM-specific SARS-CoV neutralizing mAbs can act as ACE2 mimics triggering the fusogenic rearrangmement of the S protein^36^. We evaluated mAbs of different epitope specificity (**Extended Data Table 1**) to induce fusogenic rearrangement of soluble S trimers as measured by negative stain electron microscopy imaging (**Fig. 4a**). Three RBM mAbs (S2E12, S2X58, and S2D106) triggered rearrangement to the postfusion state of a native SARS-CoV-2 S ectodomain trimer, likely due to conformational selection for open RBDs. S2E12 and S2X58 triggered a rapid rearrangement of S, whereas S2D106 did so more slowly. As expected, another RBM mAb (S2M11) that locks neighbouring RBDs in a closed state^5^ did not induce fusogenic S rearrangements. Antibodies to NTD (S2X333) and to the proteoglycan site at the base of RBD (S309) also did not trigger rearrangement due to the absence of conformational selection for open RBDs.

**Fig. 4.**
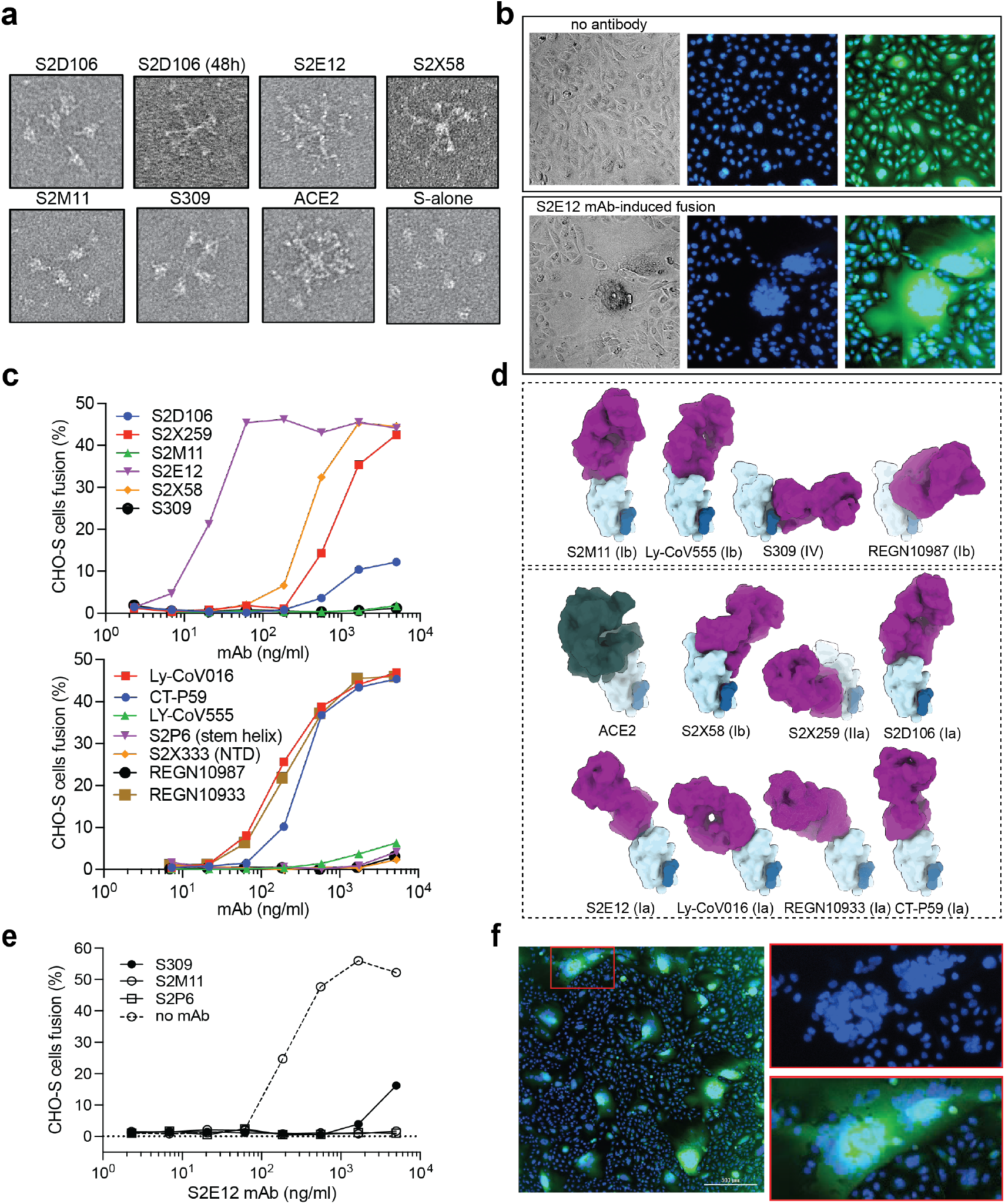
RBM mAbs trigger the fusogenic rearrangmement of the S protein and promote membrane fusion. **a,** MAbs or soluble ACE2 were incubated for 1 hour with native-like soluble prefusion S trimer of SARS-CoV-2 to track by negative stain EM imaging the fusogenic rearrangement of soluble S trimers visible as rosettes. **b**, Cell-cell fusion of CHO cells expressing SARS-CoV-2 S (CHO-S) on the plasma membrane in the absence (upper panel) or presence of 5 μg/ml of S2E12 mAb (lower panel) as detected by immuno-fluorescence. Nuclei stained with Hoechst dye; cytoplasm stained with CellTracker Green. (**c**), CHO-S cell-cell fusion mediated by different spike-specific mAbs quantified as described in Methods. **d**, Structures of 11 Fab-RBD complexes related to mAbs used in (c) (RBD orientation is fixed) and of ACE2-RBD as determined by a combination of X-ray crystallography and cryo-EM analysis (PDBs, **Extended Data Table 1**). Shown in parentheses the RBD antigenic site as defined according to Piccoli et al.^3^ **e**, Inhibition of S2E12-induced cell-cell fusion performed as in (c) by a fixed amount (15 μg/ml) of indicated mAbs. **f**, Trans-fusion of S-positive CHO cells with S-negative fluorescently-labelled CHO cells. Staining as in (b).

To investigate whether the antibody-mediated triggering of fusogenic rearrangement could promote membrane fusion, we evaluated a panel of mAbs for their capacity to induce cell-cell fusion of CHO cells (lacking ACE2 expression) stably transduced with full-length SARS-CoV-2 S (**Fig. 4b**). Syncytia formation was triggered by all mAbs recognizing antigenic sites Ia and IIa (**Extended Data Table 1**), that are only accessible in the open RBD state, with EC_50_ values ranging from 20 ng/ml for S2E12 to >1 μg/ml for S2D106 (**Fig. 4c-d**). Syncytia were also formed by the three clinical-stage mAbs REGN10933 (casirivimab), Ly-CoV016 (etesevimab) and CT-P59 (regdanvimab). In contrast, syncytia were not formed in the presence of mAbs binding to the open and closed RBD states (S2M11, S309, bamlanivimab and imdevimab), to the NTD (S2X333) and to a conserved site in the S2 subunit stem helix (S2P6). An interesting exception is provided by S2X58, a mAb that was structurally defined in this study as binding to the site Ib, which is accessible on open and closed RBDs (**Extended Data Fig. 6).** However, due to steric clashes between the S2X58 Fab and the NTD of a neighboring monomer in the closed S state, this mAb appears to conformationally select the open RBDs, thus explaining its fusogenic activity. With regard to the possible interaction between fusogenic and non-fusogenic antibodies, we found that syncytia formation induced by S2E12 could be inhibited by different classes of antibodies comprising S2M11 (that locks RBDs in a closed state), S309 (targeting a proteoglycan site at the base of RBD) and S2P6 (destabilizing the stem helix in S2) (**Fig. 4e**). These results highlight that different combinations of antibodies may interfere with each other by promoting or inhibiting membrane fusion.

To address if antibodies may promote cell-to-cell spread of the infection, we co-cultured S-positive CHO cells with S-negative fluorescently labelled CHO cells. In these conditions S2E12 mAb promoted unidirectional fusion of S-positive CHO cells with S-negative CHO cells in the absence of ACE2 (defined here as “trans-fusion”) (**Fig. 4f**). To address if this mechanism may also mediate ACE2 independent infection of tethered virus, we infected HeLa-DC-SIGN cells with live SARS-CoV-2-Nluc virus in the presence of fusion-enhancing mAbs at different dilutions. In these conditions S2E12, S2D106 and S2X58 failed to promote infection (**Extended Data Fig. 7**). Collectively, these findings indicate that in certain conditions of antibody concentration and cell-to-cell proximity a sub-class of RBM antibodies that select the open conformation of RBD may promote cell-to-cell fusion with ACE2-negative cells. However, the fusogenic activity of these mAbs may not be sufficient to promote entry of virions tethered to the cell surface in the absence of ACE2. It remains to be established whether under other conditions RBM mAbs may mediate ACE2-independent SARS-CoV-2 entry, as previously observed for anti-MERS-CoV neutralizing mAbs captured by FcγRIIa expressing cells in vitro^37^.

### Membrane lectin receptors modulate neutralizing activity by different classes of antibodies

Given the dual function of certain RBM antibodies in inhibiting ACE2 binding and triggering fusion, and the dependence on auxiliary receptor expression of neutralization by specific antibodies, we compared the neutralizing activity of a panel of mAbs using authentic SARS-CoV-2 and target cells expressing different levels of ACE2 and lectin receptors. When tested on cells over-expressing ACE2, all anti-RBM mAbs (S2E12, S2D106, S2X58 and S2M11) potently neutralized infection, while non-RBM mAbs S309 and S2X333 failed to do so (**Fig. 2** and **Fig. 5a**). However, when tested on cells expressing low levels of ACE2 together with SIGLEC1, DC-SIGN or L-SIGN, S309 and S2X333 showed enhanced neutralizing activity, with S309 reaching 100% of neutralization (**Fig. 5b-d** and **Extended Data Fig. 8**). Intriguingly, while all RBM mAbs retained neutralizing activity on SIGLEC1^+^ cells, two of the RBM mAbs (S2D106 and S2X58) lost neutralizing activity on cells expressing DC-SIGN or L-SIGN showing only partial neutralization at the highest concentrations tested. Of note, the S2E12 RBM mAb retained substantial neutralizing activity on lectin expressing cells. The only RBM mAb that retained potent neutralizing activity on all target cells was S2M11, consistent with its capacity to lock the RBDs in the closed state^5^, thereby preventing access of RBM-specific antibodies to their cognate epitope. The loss of neutralizing activity of S2X58 and S2D106 mAbs observed on DC-SIGN and L-SIGN expressing cells was confirmed with both live SARS-CoV-2 (wildtype), as well as with live SARS-CoV-2-Nluc (**Extended Data Fig. 8**). Collectively these data show that mAbs that do not block ACE2 attachment, such as S309 or S2X333, potently neutralize ACE2-dependent, lectin-facilitated infection of target cells, highlighting that they may act by inhibiting steps in viral binding and entry independent on the interaction between the RBM and ACE2 (**Fig. 5e**).

**Fig. 5.**
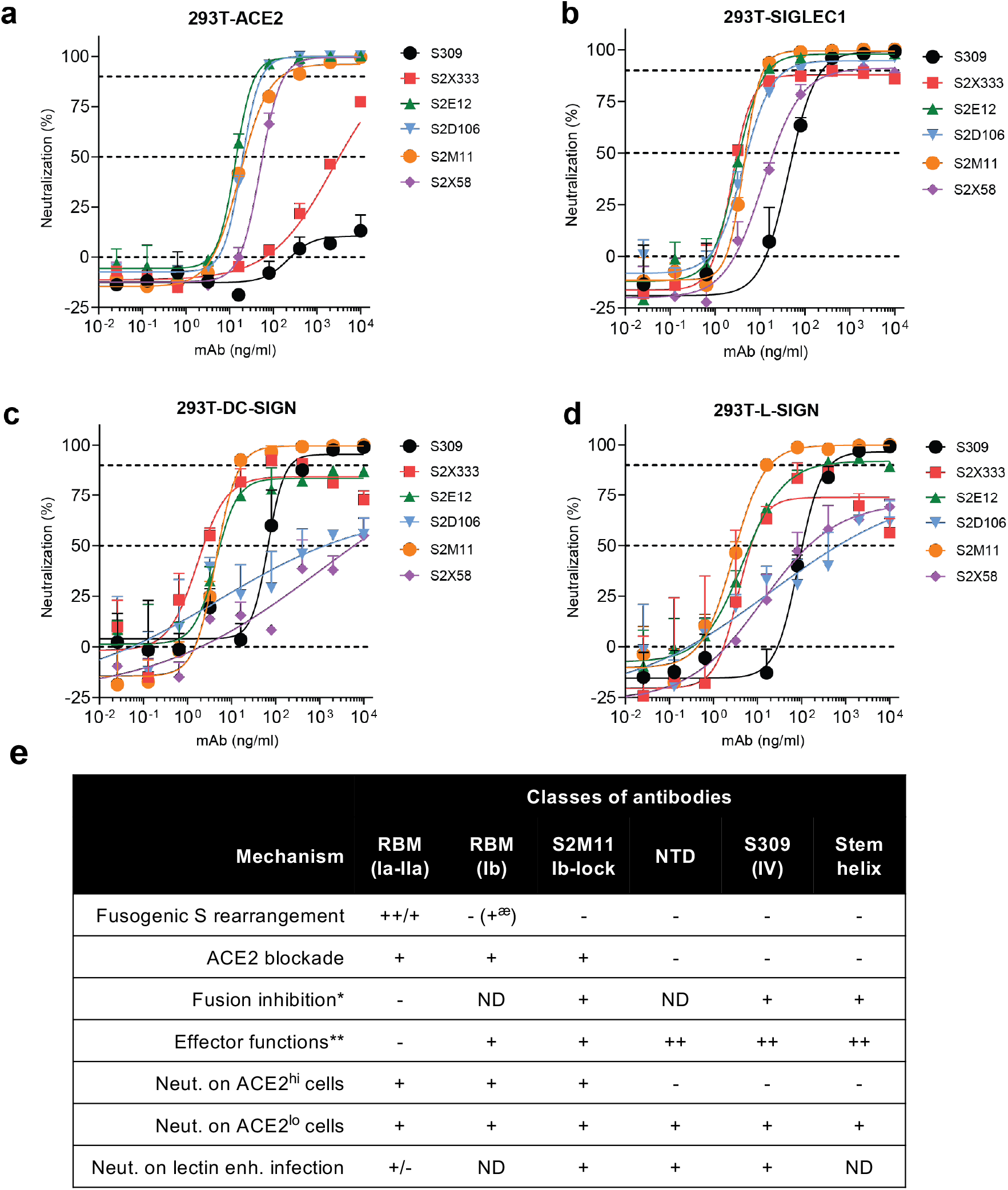
SIGLEC1, DC-SIGN and L-SIGN modulate neutralizing activity by different classes of antibodies. **a-d**, Neutralization of infection by authentic SARS-CoV-2 pre-incubated with indicated mAbs of HEK293T cell lines stably overexpressing DC-SIGN, L-SIGN, SIGLEC1 or ACE2. Infection was measured by immunostaining at 24 hours for the SARS-CoV-2 nucleoprotein. **e**, Summary of the mechanisms of action of different classes of spike-specific mAbs based on this and previous studies. *, mAb-mediated inhibition of fusion between CHO-spike cells and ACE2+ Vero-E6 cells; **, based on mAb-dependent activation of human FcγRs performed with a bioluminescent reporter assay as in^27^. ^æ^, S2X58 binds to open RDB due to a confomational clash with neighboring NTD

## Discussion

We have shown that transmembrane lectins act as auxiliary receptors, rather than entry receptors for SARS-CoV-2^14,15^, thus facilitating the infection via the canonical ACE2 pathway. This finding likely addresses the efficiency of lower respiratory tract infection despite the paradoxically low level of ACE2 expression, even in the presence of interferon^38,39^ which induces the production of an ACE2 isoform that does not bind to SARS-CoV-2 S^40,41^. The auxiliary role of lectins in SARS-CoV-2 infection is in line with the known biology of these adhesion molecules that bind glycans characteristic of cellular membranes and pathogen surfaces to promote trans-infection^42^. Among the three auxiliary receptors, SIGLEC1 is of particular relevance because this receptor is prominently expressed in alveolar macrophages in association with viral RNA, thus supporting a model of trans-infection, tissue dissemination and the triggering of immune responses by myeloid cells, rather than a direct target for productive infection^43^. DC-SIGN and L-SIGN association with SARS-CoV-2 is also of relevance because of their interaction with blood endothelium and role in immune activation.

Expression of lectin receptors can also influence the neutralizing activity of different classess of spike-specific monoclonal antibodies, In addition, the present work identifies the the ability of various mAbs to interfere with fusion events. We expand our initial observation on SARS-CoV and MERS-CoV^36,37^ by showing that most RBM mAbs can trigger the fusogenic rearrangement of S, albeit with varying efficiency. By stabilizing the RBDs in the open conformations, these antibodies might act as receptor mimics. This finding suggests that premature conformational triggering resulting in loss of the potential of a spike protein to engender productive infection - we term this mechanism spike inactivation herein - may be the prominent mode of viral neutralization for this class of antibodies, over the simple inhibition of receptor binding. However, we have also shown that these antibodies can promote fusion of Spike-expressing cells with neighboring cells, even if the latter lack ACE2. Intriguingly, the formation of syncytia has been observed in autopsy samples from severe COVID-19 cases^35,44–46^. It is tempting to speculate that fusogenic antibodies although highly effective in preventing and controlling the early phases of infection^7,8,47^, may contribute at a later stage to the spread of infection and inflammation.

Overall, our study highlights the novel finding that ranking of SARS-CoV-2 neutralizing antibodies is highly dependent on the level of ACE2 expression and on the presence of auxiliary receptors and identifies a mechanism that might possibly result in creation of multinucleate viral factories potentially enhanced by specific antibodies. Such antibodies can be protective in vivo regardless of the capacity to trigger, but this protective capacity may represent a balance between neutralization and induction of fusion between infected cells in vivo. Our results suggest that assessment of both vaccines and mAbs needs to include their effects on steps in viral infection identified in this study.

## Acknowledgements

We thank Nuria Izquierdo-Useros and Javier Martinez-Picado for useful commentary.

## Author contributions

Conceived study: F.A.L., L.S., A.L., L.P., D.V., A.T. and D.C. Designed study and experiments: F.A.L., L.S., F.B. and D.C. Performed in vitro virological experiments: F.A.L., F.B., Y-J.P., S.B., M.M-R., J.N., A.C.W., J.E.B., J.Z., H.K, M.A. EM data collection and analysis: Y-J.P., A.C.W., J.E.B. Produced antibodies for in vitro and in vivo studies: S.J. and E.C. Recombinant glycoprotein production: J.E.B., M.M., E.D. Hamster model and data analysis: F.B. Bioinformatic analysis: L.S., A.T. Manuscript writing: F.A.L, F.B., L.P., D.V., A.L., A.T. and D.C. Supervision: L.P., D.V., H.W.V, A.T. and D.C.

## MATERIALS AND METHODS

### Generation of stable overexpression cell lines

Lentiviruses were generated by co-transfection of Lenti-X 293T cells (Takara) with lentiviral expression plasmids encoding DC-SIGN (CD209), L-SIGN (CLEC4M), SIGLEC1, TMPRSS2 or ACE2 (all obtained from Genecopoeia) and the respective lentiviral helper plasmids. Forty-eight hours post transfection, lentivirus in the supernatant was harvested and concentrated by ultracentrifugation for 2 h at 20,000 rpm. Lenti-X 293T (Takara), Vero E6 (ATCC), MRC5 (Sigma-Aldrich), A549 (ATCC) or HeLa (ATCC) cells were transduced in the presence of 6 ug/mL polybrene (Millipore) for 24 h. Cell lines overexpressing two transgenes were transduced subsequently. Selection with puromycin and/or blasticidin (Gibco) was started two days after transduction and selection reagent was kept in the growth medium for all subsequent culturing. Single cell clones were derived from the A549-ACE2-TMPRSS2 cell line, all other cell lines represent cell pools.

### SARS-CoV-2 neutralization

Cells cultured in DMEM supplemented with 10% FBS (VWR) and 1x Penicillin/Streptomycin (Thermo Fisher Scientific) were seeded in black 96-well plates at 20,000 cells/well. Serial 1:4 dilutions of the monoclonal antibodies were incubated with 200 pfu of SARS-CoV-2 (isolate USA-WA1/2020, passage 3, passaged in Vero E6 cells) for 30 min at 37°C in a BSL-3 facility. Cell supernatant was removed and the virus-antibody mixture was added to the cells. 24 h post infection, cells were fixed with 4% paraformaldehyde for 30 min, followed by two PBS (pH 7.4) washes and permeabilization with 0.25% Triton X-100 in PBS for 30 min. After blocking in 5% milk powder/PBS for 30 min, cells were incubated with a primary antibody targeting SARS-CoV-2 nucleocapsid protein (Sino Biological, cat. 40143-R001) at a 1:2000 dilution for 1h. After washing and incubation with a secondary Alexa647-labeled antibody mixed with 1 ug/ml Hoechst33342 for 1 hour, plates were imaged on an automated cell-imaging reader (Cytation 5, Biotek) and nucleocapsid-positive cells were counted using the manufacturer’s supplied software.

### SARS-CoV-2-Nluc neutralization

Neutralization was determined using SARS-CoV-2-Nluc, an infectious clone of SARS-CoV-2 (based on strain 2019-nCoV/USA_WA1/2020) encoding nanoluciferase in place of the viral ORF7, which demonstrates comparable growth kinetics to wild type virus (Xie et al., Nat Comm, 2020, https://doi.org/10.1038/s41467-020-19055-7). Cells were seeded into black-walled, clear-bottom 96-well plates at 20,000 cells/well (293T cells were seeded into poly-L-lysine-coated wells at 35,000 cells/well) and cultured overnight at 37°C. The next day, 9-point 4-fold serial dilutions of antibodies were prepared in infection media (DMEM + 10% FBS). SARS-CoV-2-Nluc was diluted in infection media at the indicated MOI, added to the antibody dilutions and incubated for 30 min at 37°C. Media was removed from the cells, mAb-virus complexes were added, and cells were incubated at 37°C for 24 h. Media was removed from the cells, Nano-Glo luciferase substrate (Promega) was added according to the manufacturer’s recommendations, incubated for 10 min at RT and luciferase signal was quantified on a VICTOR Nivo plate reader (Perkin Elmer).

### SARS-CoV-2 pseudotyped VSV production and neutralization

To generate SARS-CoV-2 pseudotyped vesicular stomatitis virus, Lenti-X 293T cells (Takara) were seeded in 10-cm dishes for 80%. next day confluency. The next day, cells were transfected with a plasmid encoding for SARS-CoV-2 S-glycoprotein (YP_009724390.1) harboring a C-terminal 19 aa truncation using TransIT-Lenti (Mirus Bio) according to the manufacturer’s instructions. One day post-transfection, cells were infected with VSV(G*ΔG-luciferase) (Kerafast) at an MOI of 3 infectious units/cell. Viral inoculum was washed off after one hour and cells were incubated for another day at 37°C. The cell supernatant containing SARS-CoV-2 pseudotyped VSV was collected at day 2 post-transfection, centrifuged at 1000 x g for 5 minutes to remove cellular debris, aliquoted, and frozen at −80°C.

For viral neutralization, cells were seeded into black-walled, clear-bottom 96-well plates at 20,000 cells/well (293T cells were seeded into poly-L-lysine-coated wells at 35,000 cells/well) and cultured overnight at 37°C. The next day, 9-point 4-fold serial dilutions of antibodies were prepared in media. SARS-CoV-2 pseudotyped VSV was diluted 1:30 in media in the presence of 100 ng/mL anti-VSV-G antibody (clone 8G5F11, Absolute Antibody) and added 1:1 to each antibody dilution. Virus:antibody mixtures were incubated for 1 hour at 37°C. Media was removed from the cells and 50 μL of virus:antibody mixtures were added to the cells. One hour post-infection, 100 μL of media was added to all wells and incubated for 17-20 hours at 37°C. Media was removed and 50 μL of Bio-Glo reagent (Promega) was added to each well. The plate was shaken on a plate shaker at 300 RPM at room temperature for 15 minutes and RLUs were read on an EnSight plate reader (Perkin-Elmer).

### Transfection-based attachment receptor screen

Lenti-X 293T cells (Takara) were transfected with plasmids encoding the following receptor candidates (all purchased from Genecopoeia): ACE2 (NM_021804), DC-SIGN (NM_021155), L-SIGN (BC110614), LGALS3 (NM_002306), SIGLEC1 (NM_023068), SIGLEC3 (XM_057602), SIGLEC9 (BC035365), SIGLEC10 (NM_033130), MGL (NM_182906), MINCLE (NM_014358), CD147 (NM_198589), ASGR1 (NM_001671.4), ASGR2 (NM_080913), NRP1 (NM_003873). One day post transfection, cells were infected with SARS-CoV-2 pseudotyped VSV at 1:20 dilution in the presence of 100 ng/mL anti-VSV-G antibody (clone 8G5F11, Absolute Antibody) at 37°C. One hour post-infection, 100 μL of media was added to all wells and incubated for 17-20 hours at 37°C. Media was removed and 50 μL of Bio-Glo reagent (Promega) was added to each well. The plate was shaken on a plate shaker at 300 RPM at room temperature for 15 minutes and RLUs were read on an EnSight plate reader (Perkin-Elmer).

### Trans-infection

Parental HeLa cells or HeLa cells stably expressing DC-SIGN, L-SIGN or SIGLEC1 were seeded at 5,000 cells per well in black-walled clear-bottom 96-well plates. One day later, cells reached about 50% confluency and were inoculated with SARS-CoV-2 pseudotyped VSV at 1:10 dilution in the presence of 100 ng/mL anti-VSV-G antibody (clone 8G5F11, Absolute Antibody) at 37°C for 2 h. For antibody-mediated inhibition of trans-infection, cells were pre-incubated with 10 ug/mL anti-SIGLEC1 antibody (Biolegend, clone 7-239) for 30 min. After 2 h inoculation, cells were washed four times with complete medium and 10,000 VeroE6-TMPRSS2 cells per well were added and incubated 17-20 h at 37°C for trans-infection. Media was removed and 50 μL of Bio-Glo reagent (Promega) was added to each well. The plate was shaken on a plate shaker at 300 RPM at room temperature for 15 minutes and RLUs were read on an EnSight plate reader (Perkin-Elmer).

### Cell-cell fusion of CHO-S cells

CHO cells stably expressing SARS-CoV-2 S-glycoprotein were seeded in 96 well plates for microscopy (Thermo Fisher Scientific) at 12’500 cells/well and the following day, different concentrations of mAbs and nuclei marker Hoechst (final dilution 1:1000) were added to the cells and incubated for additional 24h hours. Fusion degree was established using the Cytation 5 Imager (BioTek) and an object detection protocol was used to detect nuclei as objects and measure their size. The nuclei of fused cells (i.e., syncytia) are found aggregated at the center of the syncitia and are recognized as a unique large object that is gated according to its size. The area of the objects in fused cells divided by the total area of all the object multiplied by 100 provides the percentage of fused cells

### Negative stain EM imaging the fusogenic rearrangement of soluble S trimers

SARS-CoV-2 S ectodomain trimer was engineered as follow and recombinantly expressed. The D614G SARS-CoV-2 S has a mu-phosphatase signal peptide beginning at 14Q, a mutated S1/S2 cleavage site (SGAR), ends at residue 1211K and followed by a TEV cleavage, fold-on trimerization motif, and an 8X his tag in the pCMV vector. 10 μM S was incubated with 13uM Fab/protein for 1 or 48 hours at room temperature. Samples were diluted to be 0.01 mg/mL immediately before protein was adsorbed to glow-discharged carbon-coated copper grids for ~30seconds prior to a 2% uranyl formate staining. Micrographs were recorded using the Leginon software^48^ on a 100kV FEI Tecnai G2 Spirit with a Gatan Ultrascan 4000 4k x 4k CCD camera at 67,000 nominal magnification. The defocus ranged from 1.0 to 2.0 μm and the pixel size was 1.6 Å.

### Cryo-electron microscopy

SARS-CoV-2 HexaPro S^49^ at 1.2 mg/mL was incubated with 1.2 fold molar excess of recombinantly purified S2X58 for 10 seconds at room temperature before application onto a freshly glow discharged 2.0/2.0 UltrAuFoil grid (200 mesh). Plunge freezing used a vitrobot MarkIV (Thermo Fisher Scientific) using a blot force of 0 and 6.5 second blot time at 100% humidity and 23°C.

Data were acquired using an FEI Titan Krios transmission electron microscope operated at 300 kV and equipped with a Gatan K2 Summit direct detector and Gatan Quantum GIF energy filter, operated in zero-loss mode with a slit width of 20 eV. Automated data collection was carried out using Leginon^48^ at a nominal magnification of 130,000x with a pixel size of 0.525 Å and stage tilt angles up to 35°. The dose rate was adjusted to 8 counts/pixel/s, and each movie was acquired in super-resolution mode fractionated in 50 frames of 200 ms. 4,126 micrographs were collected with a defocus range between −0.5 and −3.0 μm. Movie frame alignment, estimation of the microscope contrast-transfer function parameters, particle picking, and extraction were carried out using Warp^50^. Particle images were extracted with a box size of 800 binned to 400 pixels2 yielding a pixel size of 1.05 Å.

Two rounds of reference-free 2D classification were performed using CryoSPARC^51^ to select well-defined particle images. These selected particles were subjected to two rounds of 3D classification with 50 iterations each (angular sampling 7.5° for 25 iterations and 1.8° with local search for 25 iterations), using our previously reported closed SARS-CoV-2 S structure as initial model (PDB 6VXX) in Relion^52^. 3D refinements were carried out using non-uniform refinement along with per-particle defocus refinement in CryoSPARC^53^. Selected particle images were subjected to the Bayesian polishing procedure implemented in Relion3.0^54^ before performing another round of non-uniform refinement in CryoSPARC followed by per-particle defocus refinement and again non-uniform refinement. Local resolution estimation, filtering, and sharpening were carried out using CryoSPARC. Reported resolutions are based on the gold-standard Fourier shell correlation (FSC) of 0.143 criterion and Fourier shell correlation curves were corrected for the effects of soft masking by high-resolution noise substitution. UCSF ChimeraX^55^ and Coot^56^ were used to fit atomic models into the cryoEM maps.

### Immunofluorescence analysis

HEK293T-derived cell lines were seeded onto poly-D-Lysine-coated 96-well plates (Sigma-Aldrich) and fixed 24 h after seeding with 4% paraformaldehyde for 30 min, followed by two PBS (pH 7.4) washes and permeabilization with 0.25% Triton X-100 in PBS for 30 min. Cells were incubated with primary antibodies anti-DC-SIGN/L-SIGN (Biolegend, cat. 845002, 1:500 dilution), anti-DC-SIGN (Cell Signaling, cat. 13193S, 1:500 dilution), anti-SIGLEC1 (Biolegend, cat. 346002, 1:500 dilution) or anti-ACE2 (R&D Systems, cat. AF933, 1:200 dilution) diluted in 3% milk powder/PBS for 2 h at room temperature. After washing and incubation with a secondary Alexa647-labeled antibody mixed with 1 ug/ml Hoechst33342 for 1 hour, plates were imaged on an inverted fluorescence microscope (Echo Revolve).

### ACE2/TMPRSS2 RT-qPCR

RNA was extracted from the cells using the NucleoSpin RNA Plus kit (Macherey-Nagel) according to the manufacturer’s protocol. RNA was reverse transcribed using the High Capacity cDNA Reverse Transcription kit (Applied Biosystems) according to the manufacturer’s instructions. Intracellular levels of ACE2 (Forward Primer: CAAGAGCAAACGGTTGAACAC, Reverse Primer: CCAGAGCCTCTCATTGTAGTCT), HPRT (Forward Primer: CCTGGCGTCGTGATTAGTG, Reverse Primer: ACACCCTTTCCAAATCCTCAG), and TMPRSS2 (Forward Primer: CAAGTGCTCCRACTCTGGGAT, Reverse Primer: AACACACCGRTTCTCGTCCTC) were quantified using the Luna Universal qPCR Master Mix (New England Biolabs) according to the manufacturer’s protocol. Levels of ACE2 and TMPRSS2 were normalized to HPRT. Hela cells were used as the reference sample. All qPCRs were run on a QuantStudio 3 Real-Time PCR System (Applied Biosystems).

### SARS2 D614G Spike Production and biotinylation

Prefusion-stabilized SARS2 D614G spike (comprising amino acid sequence Q14 to K1211) with a C-terminal TEV cleavage site, T4 bacteriophage fibritin foldon, 8x His-, Avi- and EPEA-tag was transfected into HEK293 Freestyle cells, using 293fectin as a transfection reagent. Cells were left to produce protein for three days at 37°C. Afterwards, supernatant was harvested by centrifuging cells for 30 minutes at 500 xg, followed by another spin for 30 minutes at 4000 xg. Cell culture supernatant was filtered through a 0.2 um filter and loaded onto a 5 mL C-tag affinity matrix column, pre-equilibrated with 50 mM Tris pH 8 and 200 mM NaCl. SARS2 D614G spike was eluted, using 10 column volumes of 100 mM Tris, 200 mM NaCl and 3.8 mM SEPEA peptide. Elution peak was concentrated and injected on a Superose 6 increase 10/300 GL gel filtration column, using 50 mM Tris pH 8 and 200 mM NaCl as a running buffer. SEC fractions corresponding to monodisperse SARS2 D614G spike were collected and flash frozen in liquid nitrogen for storage at −80°C. Purified SARS2 D614G spike protein was biotinylated using BirA500 biotinylation kit from Avidity. To 50 ug of spike protein, 5 ug of BirA, and 11 uL of BiomixA and BiomixB was added. Final spike protein concentration during the biotinylation reaction was ~1 uM. The reaction was left to proceed for 16 hours at 4°C. Then, protein was desalted using two Zeba spin columns pre-equilibrated with 1x PBS pH 7.4.

### Flow cytometry analysis for DC-SIGN, L-SIGN, SIGLEC1 and ACE-2

HEK293T cells expressing DC-SIGN, L-SIGN, SIGLEC1 or ACE2 were resuspended at 4×10^6^ cells/mL and 100 μL per well were seeded onto V-bottom 96-well plates (Corning, 3894). The plate was centrifuged at 2,000 rpm for 5 minutes and washed with PBS (pH 7.4). The cells were resuspended in 200 μL of PBS containing Ghost violet 510 viability dye (Cell Signaling, cat. 13-0870-T100, 1:1,000 dilution), incubated for 15 minutes on ice and then washed. The cells were resuspended in 100 μL of FACS buffer prepared with 0.5% BSA (Sigma-Aldrich) in PBS containing the primary antibodies at a 1:100 dilution: mouse anti-DC/L-SIGN (Biolegend, cat. 845002), rabbit anti-DC-SIGN (Cell Signaling, cat. 13193), mouse anti-SIGLEC1 (Biologend, cat. 346002) or goat anti-ACE2 (R&D Systems, cat. AF933). After 1 h incubation on ice, the cells were washed two times and resuspended in FACS buffer containing the Alexa Fluor-488-labeled secondary antibodies at a 1:200 dilution: goat anti-mouse (Invitrogen cat. A11001), goat anti-rabbit (Invitrogen cat. A11008) or donkey anti-goat (Invitrogen cat. A11055). After incubation for 45 min on ice, the cells were washed three times with 200μL of FACS buffer and fixed with 200μL of 4% PFA (Alfa Aesar) for 15 mins at room temperature. Cells were washed three times, resuspended in 200μL of FACS buffer and analyzed by flow cytometry using the CytoFLEX flow cytometer (Beckman Coulter).

### Flow cytometry of SARS-CoV-2 Spike and RBD binding to cells

Biotinylated SARS-CoV-2 Spike D614G protein (Spikebiotin, in-house generated) or the biotinylated SARS-CoV-2 Spike receptor-binding domain (RBDbiotin, Sino Biological, 40592-V08B) were incubated with Alexa Fluor® 647 streptavidin (AF647-strep, Invitrogen, S21374) at a 1:20 ratio by volume for 20 min at room temperature. The labeled proteins were then stored at 4°C until further use. Cells were dissociated with TrpLE Express (Gibco, 12605-010) and 10^5^ cells were transferred to each well of a 96-well V bottom plate (Corning, 3894). Cells were washed twice in flow cytometry buffer (2% FBS in PBS (w/o Ca/Mg)) and stained with Spikebiotin-AF647-strep at a final concentration of 20 μg/ml or RBDbiotin-AF647-strep at a final concentration of 7.5 μg/ml for 1h on ice. Stained cells were washed twice with flow cytometry buffer, resuspended in 1% PFA (Electron Microscopy Sciences, 15714-S) and analyzed with the Cytoflex LX (Beckman Coulter).

### Recombinant expression of SARS-CoV-2-specific mAbs

Human mAbs were isolated from plasma cells or memory B cells of SARS-CoV-2 immune donors, as previously described ^27,57,58^. Recombinant antibodies were expressed in ExpiCHO cells at 37°C and 8% CO_2_. Cells were transfected using ExpiFectamine. Transfected cells were supplemented 1 day after transfection with ExpiCHO Feed and ExpiFectamine CHO Enhancer. Cell culture supernatant was collected eight days after transfection and filtered through a 0.2 μm filter. Recombinant antibodies were affinity purified on an ÄKTA xpress FPLC device using 5 mL HiTrap™ MabSelect™ PrismA columns followed by buffer exchange to Histidine buffer (20 mM Histidine, 8% sucrose, pH 6) using HiPrep 26/10 desalting columns

### SARS-CoV-2 infection model in hamster

#### Virus preparation

The SARS-CoV-2 strain used in this study, BetaCov/Belgium/GHB-03021/2020 (EPI ISL 109 407976|2020-02-03), was recovered from a nasopharyngeal swab taken from an RT-qPCR confirmed asymptomatic patient who returned from Wuhan, China in February 2020. A close relation with the prototypic Wuhan-Hu-1 2019-nCoV (GenBank accession 112 number MN908947.3) strain was confirmed by phylogenetic analysis. Infectious virus was isolated by serial passaging on HuH7 and Vero E6 cells ^59^; passage 6 virus was used for the study described here. The titer of the virus stock was determined by end-point dilution on Vero E6 cells by the Reed and Muench method^60^. Live virus-related work was conducted in the high-containment ABSL3 and BSL3+ facilities of the KU Leuven Rega Institute (3CAPS) under licenses AMV 30112018 SBB 219 2018 0892 and AMV 23102017 SBB 219 20170589 according to institutional guidelines.

#### Cells

Vero E6 cells (African green monkey kidney, ATCC CRL-1586) were cultured in minimal essential medium (Gibco) supplemented with 10% fetal bovine serum (Integro), 1% L-glutamine (Gibco) and 1% bicarbonate (Gibco). End-point titrations were performed with medium containing 2% fetal bovine serum instead of 10%.

#### SARS-CoV-2 infection model in hamsters

The hamster infection model of SARS-CoV-2 has been described before ^59,61^. The specific study design is shown in the schematic below. In brief, wild-type Syrian Golden hamsters (*Mesocricetus auratus*) were purchased from Janvier Laboratories and were housed per two in ventilated isolator cages (IsoCage N Biocontainment System, Tecniplast) with *ad libitum* access to food and water and cage enrichment (wood block). The animals were acclimated for 4 days prior to study start. Housing conditions and experimental procedures were approved by the ethics committee of animal experimentation of KU Leuven (license P065-2020). Female 6-8 week old hamsters were anesthetized with ketamine/xylazine/atropine and inoculated intranasally with 50 μL containing 2×10^6^ TCID50 SARS-CoV-2 (day 0).

#### Treatment regimen

Animals were prophylactically treated 48h before infection by intraperitoneal administration (i.p.) and monitored for appearance, behavior, and weight. At day 4 post infection (p.i.), hamsters were euthanized by i.p. injection of 500 μL Dolethal (200 mg/mL sodium pentobarbital, Vétoquinol SA). Lungs were collected and viral RNA and infectious virus were quantified by RT-qPCR and end-point virus titration, respectively. Blood samples were collected before infection for PK analysis.

#### SARS-CoV-2 RT-qPCR

Collected lung tissues were homogenized using bead disruption (Precellys) in 350μL RLT buffer (RNeasyMinikit, Qiagen)and centrifuged (10.000 rpm, 5 min) to pellet the cell debris. RNA was extracted according to the manufacturer’s instructions. Of 50 μL eluate, 4 μL was used as a template in RT-qPCR reactions. RT-qPCR was performed on a LightCycler96 platform (Roche) using the iTaq Universal Probes One-Step RT-qPCR kit (BioRad) with N2 primers and probes targeting the nucleocapsid^59^. Standards of SARS-CoV-2 cDNA (IDT) were used to express viral genome copies per mg tissue or per mL serum.

#### End-point virus titrations

Lung tissues were homogenized using bead disruption (Precellys) in 350 μL minimal essential medium and centrifuged (10,000 rpm, 5min, 4°C) to pellet the cell debris. To quantify infectious SARS-CoV-2 particles, endpoint titrations were performed on confluent Vero E6 cells in 96-well plates. Viral titers were calculated by the Reed and Muench method^60^ using the Lindenbach calculator and were expressed as 50% tissue culture infectious dose (TCID50) per mg tissue.

#### Histology

For histological examination, the lungs were fixed overnight in 4% formaldehyde and embedded in paraffin. Tissue sections (5 μm) were analyzed after staining with hematoxylin and eosin and scored blindly for lung damage by an expert pathologist. The scored parameters, to which a cumulative score of 1 to 3 was attributed, were the following: congestion, intra-alveolar hemorrhagic, apoptotic bodies in bronchus wall, necrotizing bronchiolitis, perivascular edema, bronchopneumonia, perivascular inflammation, peribronchial inflammation and vasculitis.

#### Binding of immunocomplexes to hamster monocytes

Immunocomplexes (IC) were generated by complexing S309 mAb (hamster IgG, either wt or N297A) with a biotinylated anti-idiotype fab fragment and Alexa-488-streptavidin, using a precise molar ratio (4:8:1, respectively). Pre-generated fluorescent IC were serially diluted incubated at 4°C for 3 hrs with freshly revitalized hamster splenocytes, obtained from a naïve animal. Cellular binding was then evaluated by cytometry upon exclusion of dead cells and physical gating on monocyte population. Results are expressed as Alexa-488 mean florescent intensity of the entire monocyte population.

#### Bioinformatic analyses

Processed Human Lung Cell Atlas (HLCA) data and cell-type annotations were downloaded from Github (https://github.com/krasnowlab/HLCA)^21^. Processed single-cell transcriptome data and annotation of lung epithelial and immune cells from SARS-CoV-2 infected individuals were downloaded from NCBI GEO database (ID: GSE158055)^22^ and Github (https://github.com/zhangzlab/covid_balf;^23^). Available sequence data from the second single-cell transcriptomics study by Liao et al^23^ were downloaded from NCBI SRA (ID: PRJNA608742) for inspection of reads corresponding to viral RNA. The proportion of sgRNA relative to genomic RNA was estimated by counting TRS-containing reads supporting a leader-TRS junction. Criteria and methods for detection of leader-TRS junction reads were adapted from Alexandersen et al.^62^. The viral genome reference and TRS annotation was based on Wuhan-Hu-1 NC_045512.2/MN908947^63^. Only 2 samples from individuals with severe COVID-19 had detectable leader-TRS junction reads (SRR11181958, SRR11181959).

---

**Extended Data Fig. 1.**
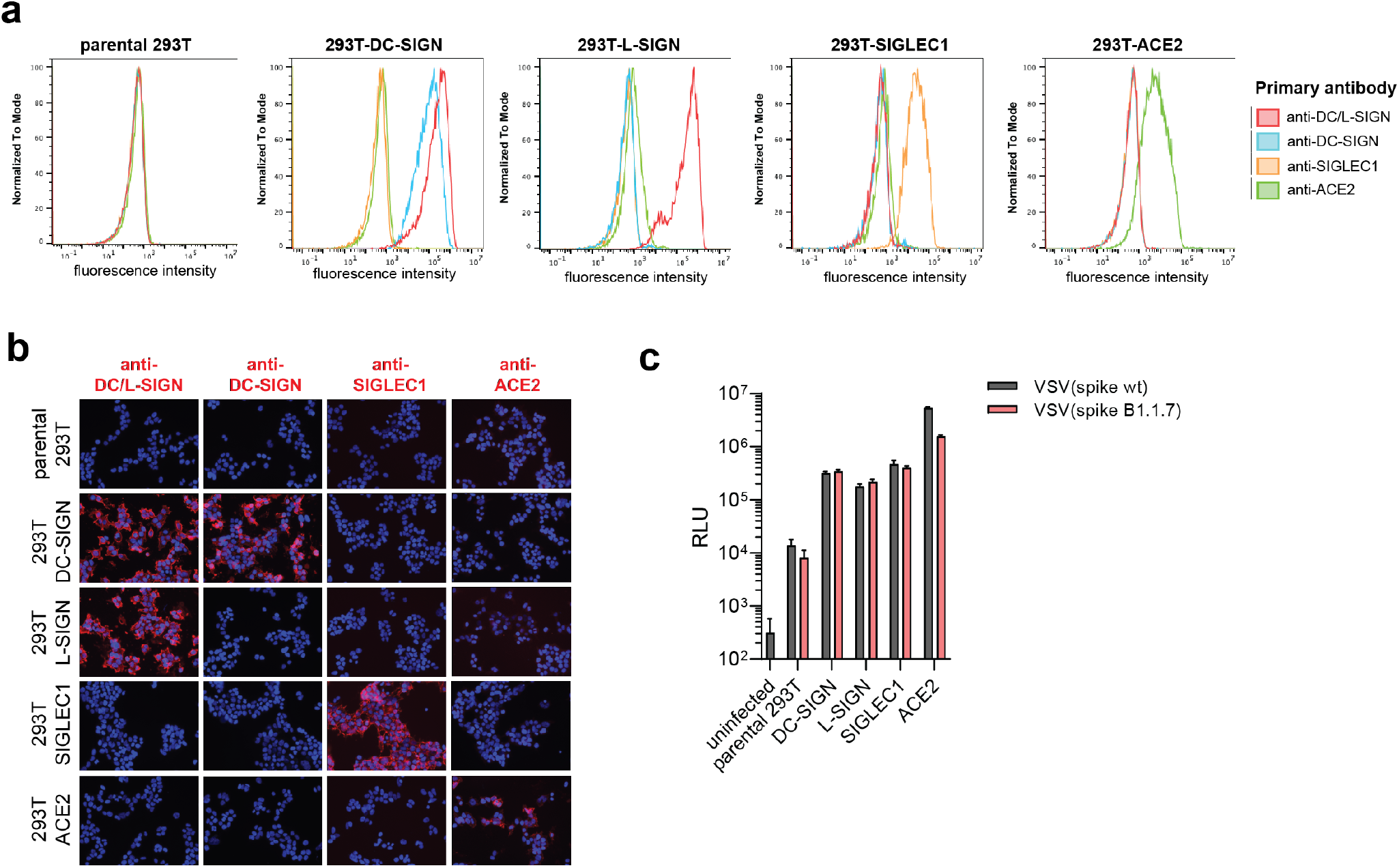
Characterization of DC-SIGN, L-SIGN and SIGLEC-1 as SARS-CoV-2 attachment factors. **a-b,** Binding of antibodies targeting DC/-L-SIGN, DC-SIGN, SIGLEC1 or ACE2 on HEK293T stably over-expressing the respective attachment receptors was analyzed by flow cytometry (a) and immunofluorescence analysis (b). **c,** HEK293T cells over-expressing the respective attachment receptors were infected with VSV-SARS-COV-2 wildtype spike (grey bars) or spike bearing mutations of the B.1.1.7 variant (red bars). Luminescence was analyzed one day post infection.

**Extended Data Fig. 2:**
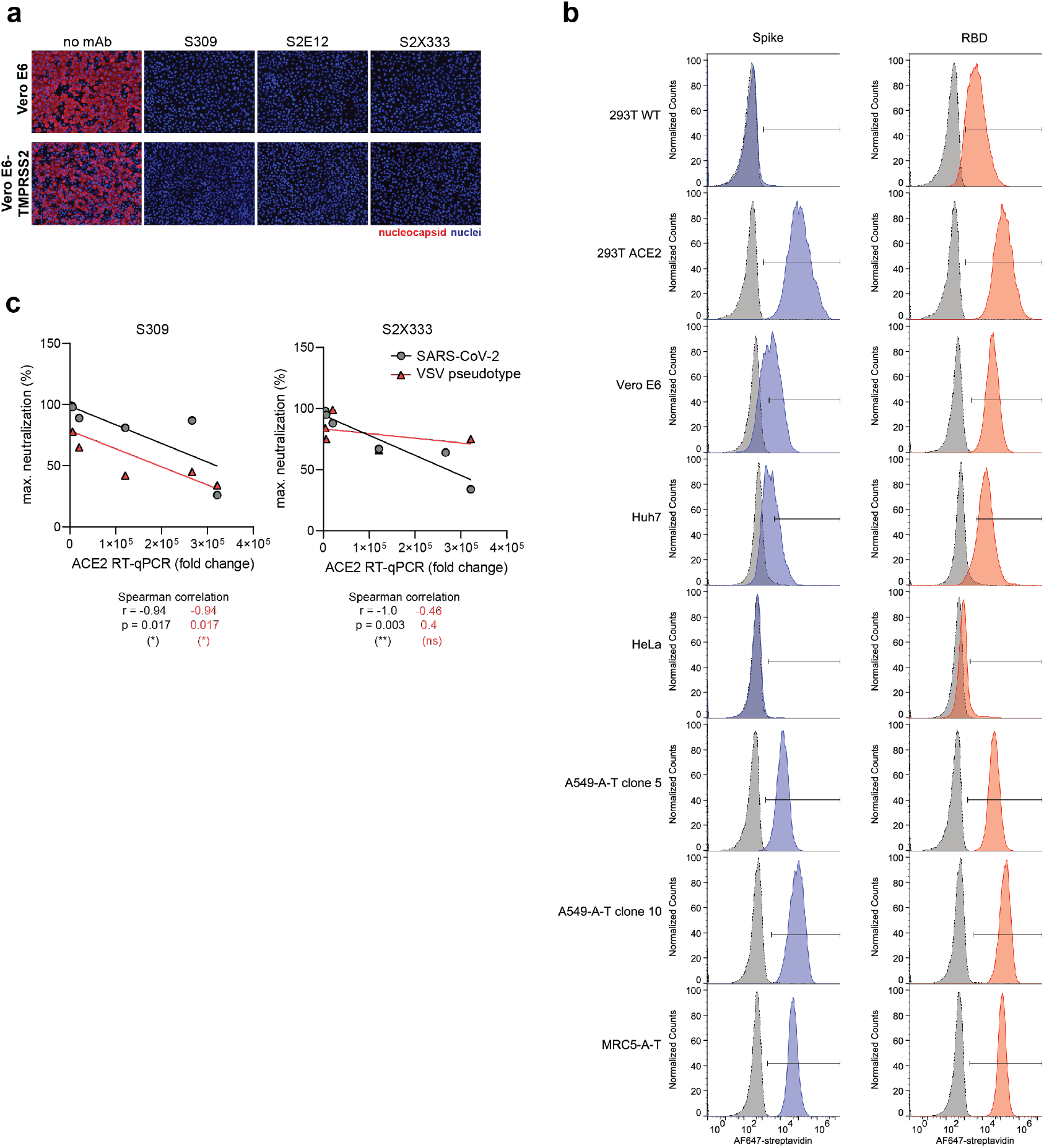
Characterization of SARS-CoV-2-susceptible cell lines. **a**, SARS-CoV-2 neutralization with 10 μg/ml of S309, S2E12 and S2X33 on Vero E6 or Vero E6-TMPRSS2 cells. Cells were infected with SARS-CoV-2 (isolate USA-WA1/2020) at MOI 0.01 in the presence of the respective mAbs. Cells were fixed 24h post infection and viral nucleocapsid protein was immunostained. **b**, Purified, fluorescently-labelled SARS-CoV-2 spike protein or RBD protein was incubated with the indicated cell lines and protein binding was quantified by flow cytometry. **c**, Correlation analysis between ACE2 transcript levels and maximum antibody neutralization in all SARS-CoV-2-susceptible cell lines.

**Extended Data Fig. 3:**
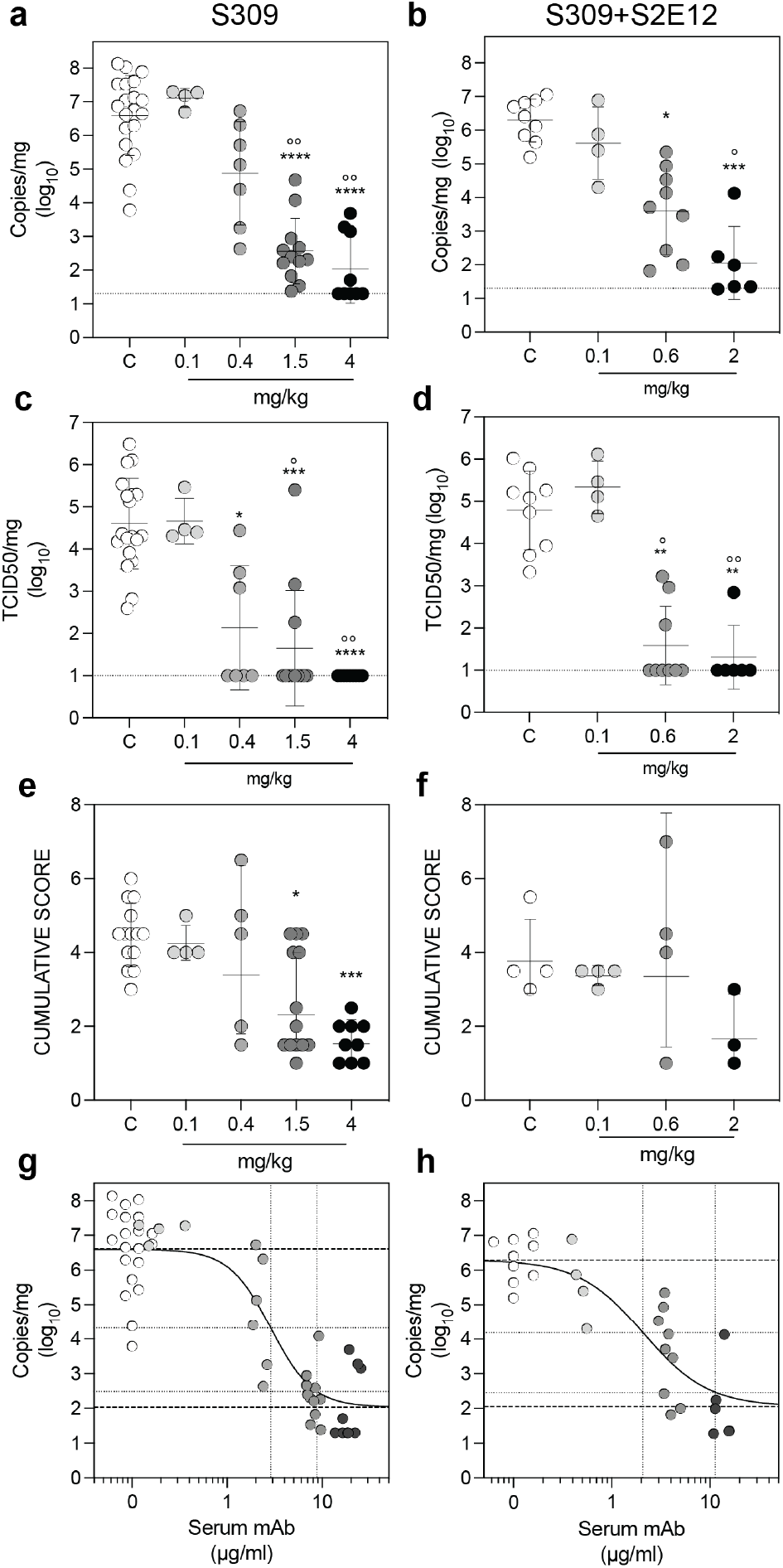
S309 or a cocktail of S309 and S2E12 provide robust in vivo protection against SARS-CoV-2 challenge. Syrian hamsters were injected with the indicated amount of mAb(s) 48 hours before intra-nasal challenge with SARS-CoV-2. (**a-b**) Quantification of viral RNA in the lungs 4 days post-infection. (**c-d**) Quantification of replicating virus in lung homogenates harvested 4 days post infection using a TCID50 assay. (**e-f**) Histopathological score of the lung tissue was assessed 4 days post infection. (**g-h**) Efficacy plots based on the correlation between the level of serum antibody measured at the time of infection and the level of SARS-CoV2 (viral RNA) measured in lungs on day 4 after infection. The dotted lines represents EC50 and EC90 for viral reduction (EC90 of S309 alone vs S309+S2E12: 9 vs 11 μg/ml, respectively).

**Extended Data Fig. 4:**
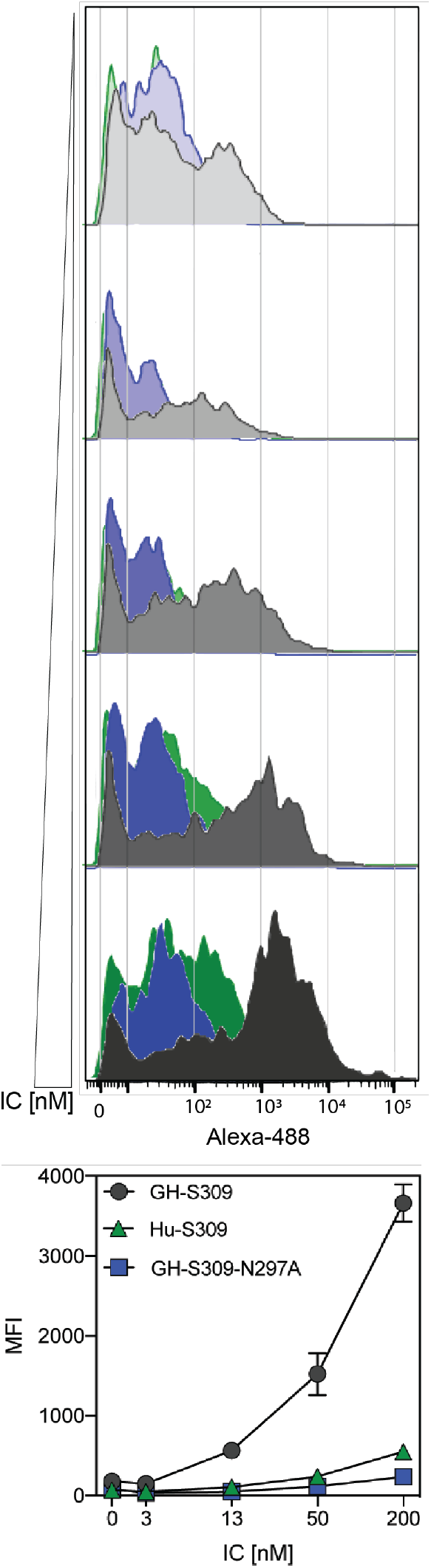
Binding of immunocomplexes to hamster splenocytes. Alexa-488 fluorescent IC were titrated (0-200 nM range) and incubated with total naïve hamster splenocytes. Binding was revealed with a cytometer upon exclusion of dead/apoptotic cells and physical gating on bona fide monocyte population. Top panel shows the fluorescent intensity associated to hamster cells of IC made with either hamster Fc antibodies (human S309, green, GH-S309, dark grey and GH-S309-N297A, blue). A single replicate of two is shown. Bottom Panel show the relative Alexa-488 mean fluorescent intensity of the replicates measured on the entire monocyte population.

**Extended Data Fig. 5:**
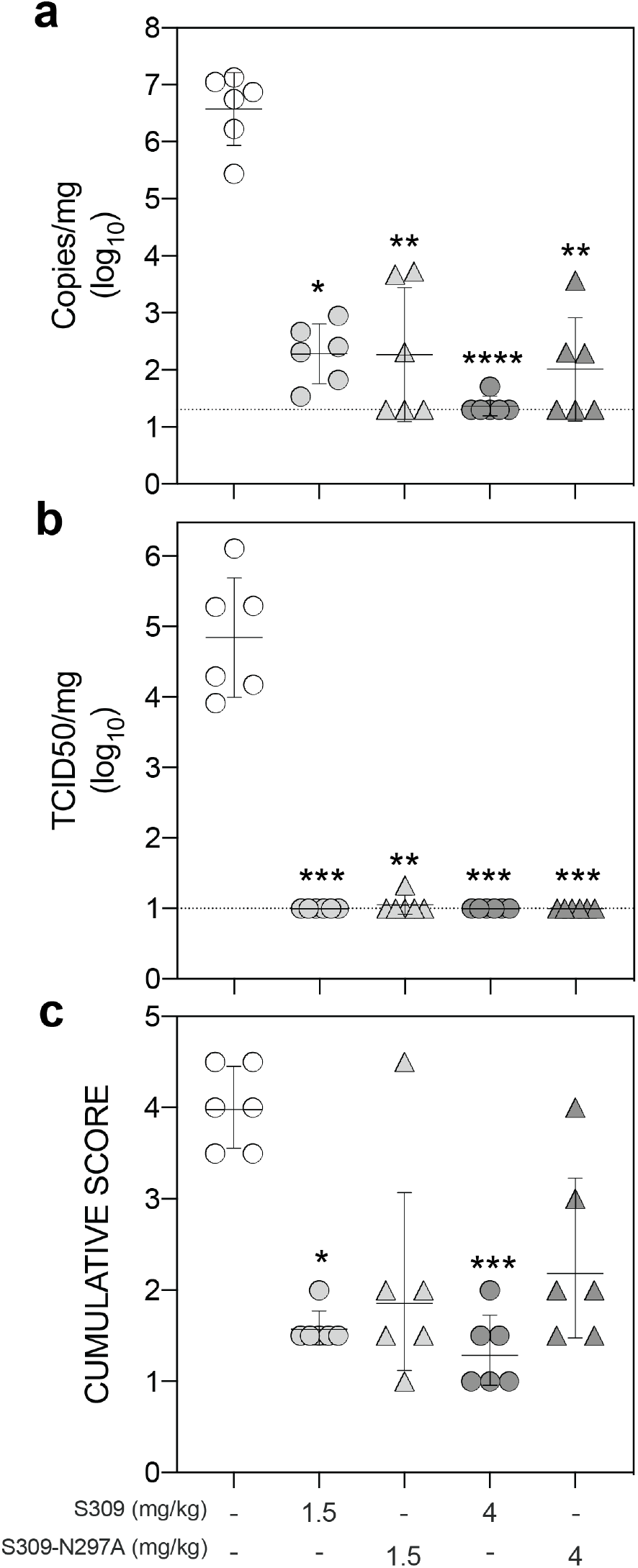
Role of host effector function in SARS-CoV-2 challenge. Syrian hamsters were injected with the indicated amount (mg/kg) of hamster IgG2a S309 either wt or Fc silenced (S309-N297A). **a**, Quantification of viral RNA in the lung 4 days post infection. **b,** Quantification of replicating virus in the lung 4 days post infection. **c,** Histopathological score in the lung 4 days post infection. Control animals (white symbols) were injected with 4 mg/kg unrelated control isotype mAb. *, **, ***, **** p< 0.05, 0.01, 0.001, 0.0001 respectively vs control animals. Mann-Whitney test.

**Extended Data Fig. 6:**
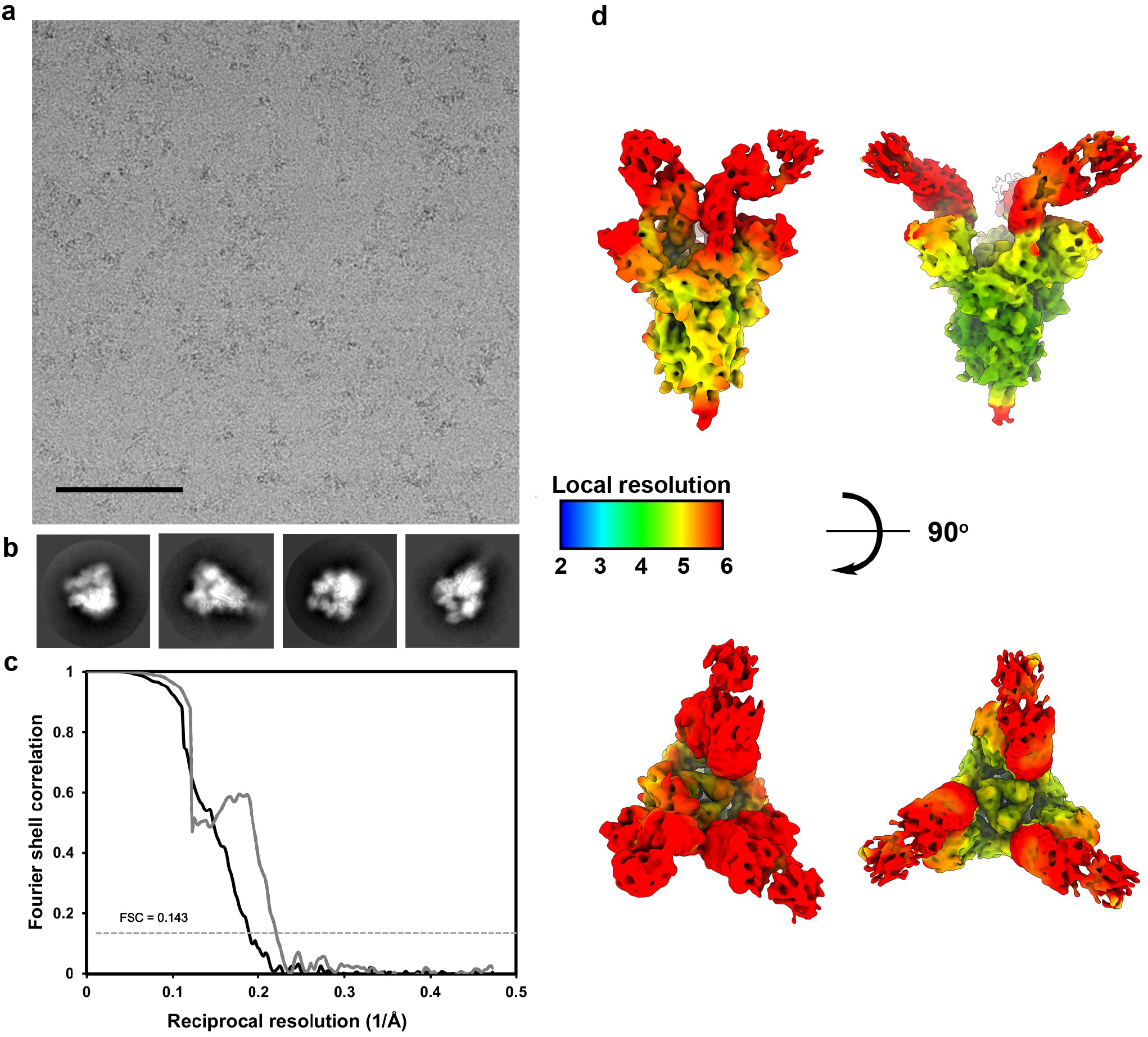
Data collection and processing of the S/S2X58 complex cryoEM datasets. **a,b**, Representative electron micrograph and 2D class averages of SARS-CoV-2 S in complex with the S2X58 Fab embedded in vitreous ice. Scale bar: 400 Å. **c**, Gold-standard Fourier shell correlation curves for the S2X58-bound SARS-CoV-2 S trimer in one RBD closed (black line) and three RBDs open conformations (gray line). The 0.143 cutoff is indicated by a horizontal dashed line. d, Local resolution maps calculated using cryoSPARC for the SARS-CoV-2 S/S2X58 Fab complex structure with one RBD closed and three RBDs open shown in two orthogonal orientations.

**Extended Data Fig. 7:**
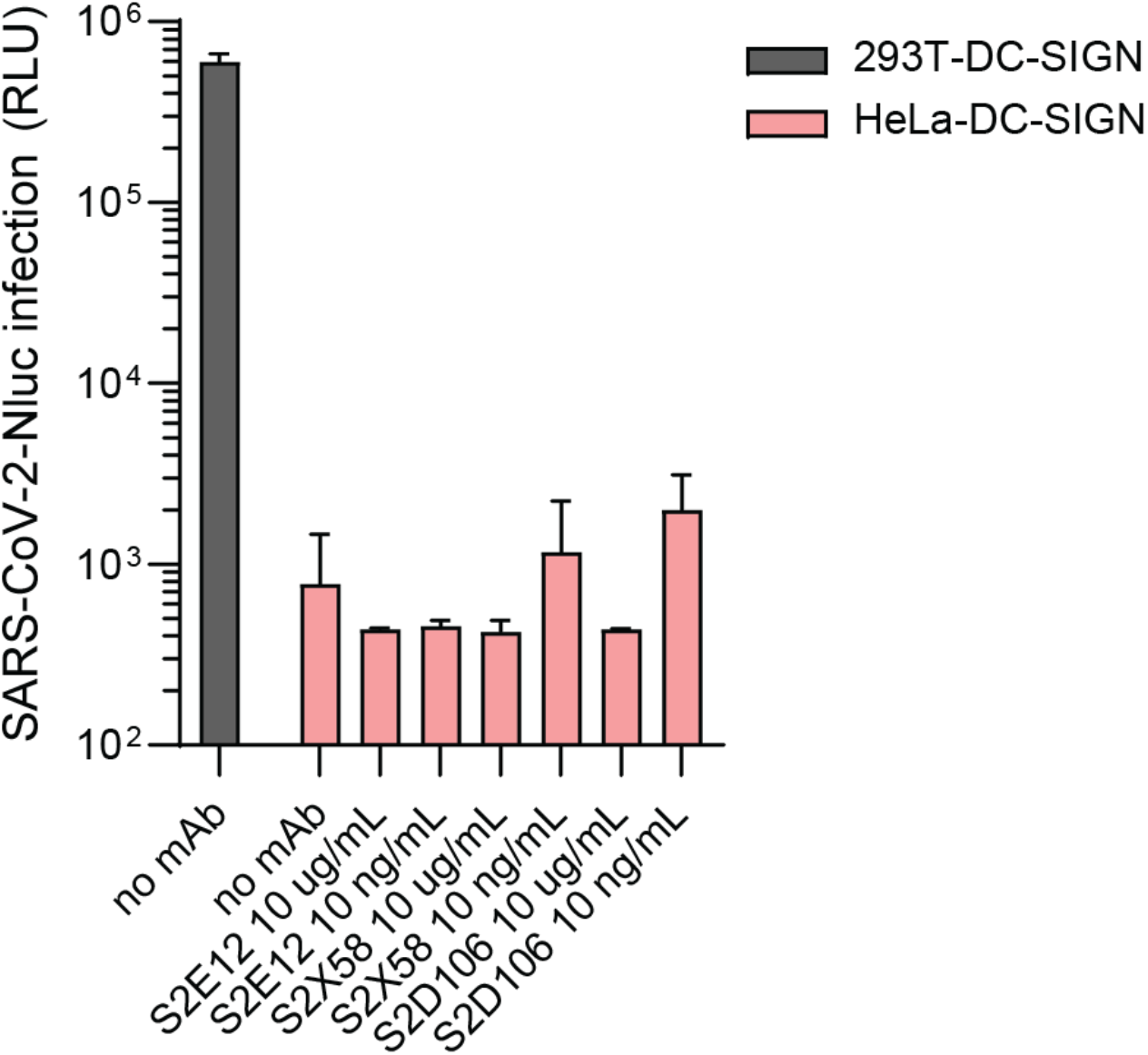
HeLa cells expressing DC-SIGN are refractory to SARS-CoV-2 infection. 293T or HeLa cells stably expressing DC-SIGN were infected with SARS-CoV-2-Nluc at MOI0.04 in the presence of the indicated antibodies. Infection was analyzed by quantification of luminescent signal at 24 h post infection.

**Extended Data Fig. 8:**
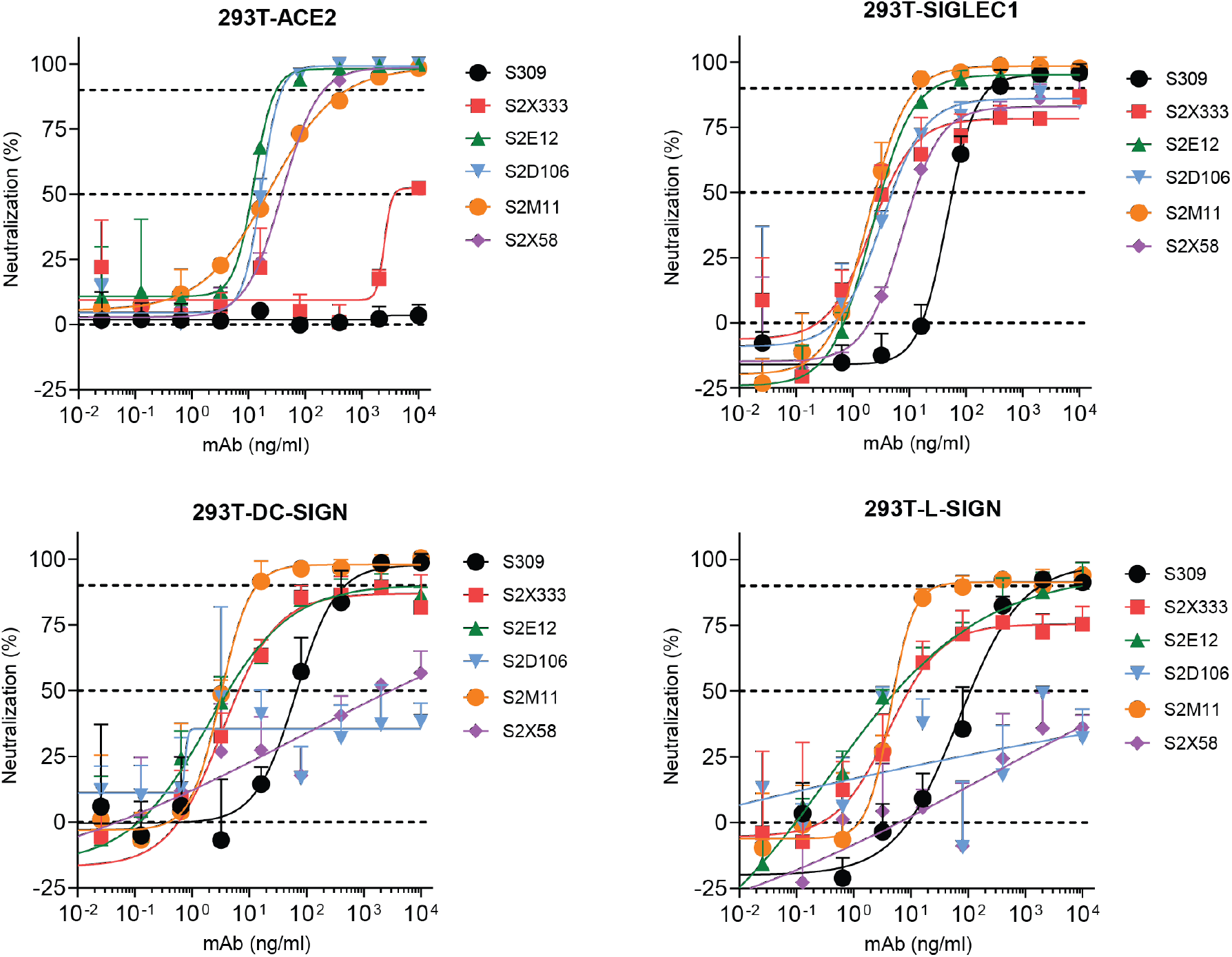
SARS-CoV-2 live virus neutralization. HEK293T cells stably expressing ACE2, SIGLEC1, DC-SIGN or L-SIGN were infected with SARS-CoV-2 at MOI 0.02 in the presence of the indicated mAbs. Cells were fixed 24h post infection, viral nucleocapsid protein was immunostained and positive cells were quantified.

**Extended Data Table 1.** MAbs used in this study.

| MAb | Alias or derived mAb(s) | Domain (site) | ACE2 blocking | SARS-CoV | Phase | PDB | Ref |
| --- | --- | --- | --- | --- | --- | --- | --- |
| S309 | VIR-7831/VIR-7832 | RBD (IV) | - | + | EUA submitted | 7IX3 | <sup>27</sup> |
| S2E12 |  | RBD (Ia) | + | - | preclinical | 7K4N | <sup>5</sup> |
| S2M11 |  | RBD (Ib-lock) | + | - | preclinical | 7K43 | <sup>5</sup> |
| S2P6 |  | S2 (stem helix) | - | + | preclinical | xxx | manuscript in preparation |
| S2X58 |  | RBD (Ib) | + | - | preclinical | this study | Starr et al. submitted |
| S2D106 |  | RBD (Ia) | + | - | preclinical | xxx | Starr et al. submitted |
| S2X259 |  | RBD (IIa) | + | - | preclinical | xxx | Tortorici et al. submitted |
| S2X333 |  | NTD (i) | - | - | preclinical | 7LXW 7LXY | <sup>28</sup> |
| Ly-CoV016 | CB6/etesevimab | RBD (Ia) | + | - | EUA in US | 7C01 | <sup>64</sup> |
| Ly-CoV555 | bamlanivimab | RBD (Ib) | + | - | EUA in US | 7KMG | <sup>65,66</sup> |
| REGN10933 | casirivimab | RBD (Ia) | + | - | EUA in US | 6XDG | <sup>6,8</sup> |
| REGN10987 | imdevimab | RBD (Ib) | + | - | EUA in US | 6XDG | <sup>6,8</sup> |
| CT-P59 | regdanvimab | RBD (Ia) | + | - | EUA in SK* | 7CM4 | <sup>67</sup> |

\*, SK, South Korea

